# Spatio-temporal mass spectrometry in living cells reveals regulators of cuproptosis

**DOI:** 10.1101/2025.07.01.662679

**Authors:** Xueliang Yang, Chong Wei, Ye Lu, Yiren Hu, Xin Liu, Gang Shao, Shuyuan Zhang, Keke Zheng, Xiaoyan Dai, Jian Kang, Chulin Sha, Qiang Zhang, Caiyun Fu

## Abstract

Cuproptosis is a unique form of regulated cell death triggered by excessive copper accumulation in mitochondria. To decipher the mechanisms of cuproptosis, we develop Spatio-Temporal Mass Spectrometry (STMS) technique to characterize dynamic subcellular proteomic changes during its early stage. STMS analysis reveals distinct subcellular proteome alterations and unique trafficking patterns over time. Through a genetic functional screen, we discover that copper stress reduces OXA1L in mitochondria, disrupting MRPL20 import and impairing oxidative phosphorylation complex I synthesis. This mitochondrial dysfunction promotes MRPL20 nuclear translocation, triggering the mitochondrial unfolded protein response as an adaptive mechanism to restore mitochondrial proteostasis during early cuproptosis. Meanwhile, AMPK phosphorylates ISCU at serine 29, stabilizing ISCU protein, and increasing the abundance of Fe-S cluster-containing proteins, which, in turn, enhances cells susceptible to cuproptosis. Building on this mechanistic insight, we develop AMPK-SPARK, a phase separation-based EGFP reporter for real-time visualization of endogenous AMPK activity in living cells during cuproptosis. Additionally, we identify COMMD4 as a cuproptosis sensor and establish a GFP-tagged COMMD4 cell model for high-content drug screening. Using this platform, we identify three FDA-approved drugs including Plerixafor, Dithranol, and Bleomycin as potential cuproptosis inhibitors. Overall, our work advances the understanding of cuproptosis regulation and provides valuable molecular tools and data resource for future research in this area.

## Main

Copper (Cu) is an essential trace element that regulates multiple biological processes critical for cell growth and function. It also serves as a catalytic cofactor for enzymes involved in energy generation and cellular metabolism^1^. Physiological Cu concentration is usually kept at a relatively low range and intricately regulated. Disrupting copper homeostasis can cause cytotoxicity and organ damages, as presented in patients with certain genetic disorders including Wilson and Menkes disease^2,3^.

Cuproptosis is a unique form of regulated cell death that was first reported in 2022. It is triggered by excessive copper accumulation within mitochondria. It is driven by ferredoxin 1 (FDX1)-mediated mitochondrial proteotoxic stress associated with protein lipoylation and degradation of iron-sulfur cluster proteins^4^. The recent studies of cuproptosis have expanded our understanding of copper metabolism and provide the opportunities to develop new treatments for copper-related diseases and cancers^5–8^. However, the complete mechanism of cuproptosis is not yet fully understood. In particular, the molecular changes involved in the early phase of cuproptosis and the cellular adaptive responses to copper stress have not yet been defined. Furthermore, the lack of sensitive and specific copper-dependent biomarkers complicates the assessment of copper dysregulation and hinders the ability to intervene in cuproptosis-related diseases.

To understand the mechanisms and identify key regulators of early cuproptosis, we exploited TurboID-mediated proximity labelling technique to characterize the spatial and temporal dynamics of the proteome among the endoplasmic reticulum (ER), Golgi, nucleus, outer mitochondrial membrane (OMM), and plasma membrane (PM) at five time points following copper influx facilitated by elesclomol (ES). We observed distinct changes in subcellular proteome and unique trafficking patterns over time. In combination with a genetic functional screen, we identified the molecules that influence cell susceptibility to cuproptosis. Further characterization identified three specific events in early cuproptosis: the translocation of MRPL20 from mitochondria into nuclei and induces mitochondrial unfolded protein response driven by OXAIL suppression, activation of AMPK-ISCU (Iron-Sulfur Cluster Assembly Enzyme) signaling, and a rapid COMMD4 protein upregulation. Building on these mechanistic insights, we developed a robust assay to accurately assess AMPK-dependent cuproptosis-related mitochondrial stress. We further conducted a high-content drug screen using COMMD4 as a cuproptosis sensor and identified the potential cuproptosis inhibitors for treating copper metabolism diseases.

### Spatial-temporal mapping of subcellular proteome dynamics in early cuproptosis

The proximity-dependent biotinylation techniques have been used to define the proteomes of different organelles in living cells^9,10^. In contrast to over 18 hrs labeling time required for BioID, TurboID and miniTurbo achieve proximity labelling in just 10 min, making them suitable for studying rapid protein dynamics^11^. To provide the spatial and temporal resolution of changes in subcellular locations during early cuproptosis, we develop a miniTurbo-based mass spectrometry technique named Spatio-Temporal Mass Spectrometry (STMS) (Fig. 1A). The biotin ligase miniTurboID was fused with a bait protein to specifically target five cellular compartments: the endoplasmic reticulum (ER), Golgi, nucleus, outer mitochondrial membrane (OMM), and plasma membrane (PM)^9^. The expression plasmids were transduced into human HT-1080 fibrosarcoma cells and the resulting V5-tagged fusion proteins were detected by Western blotting (fig S1A) and immunofluorescence (Fig. 1B).

**Fig. 1.**
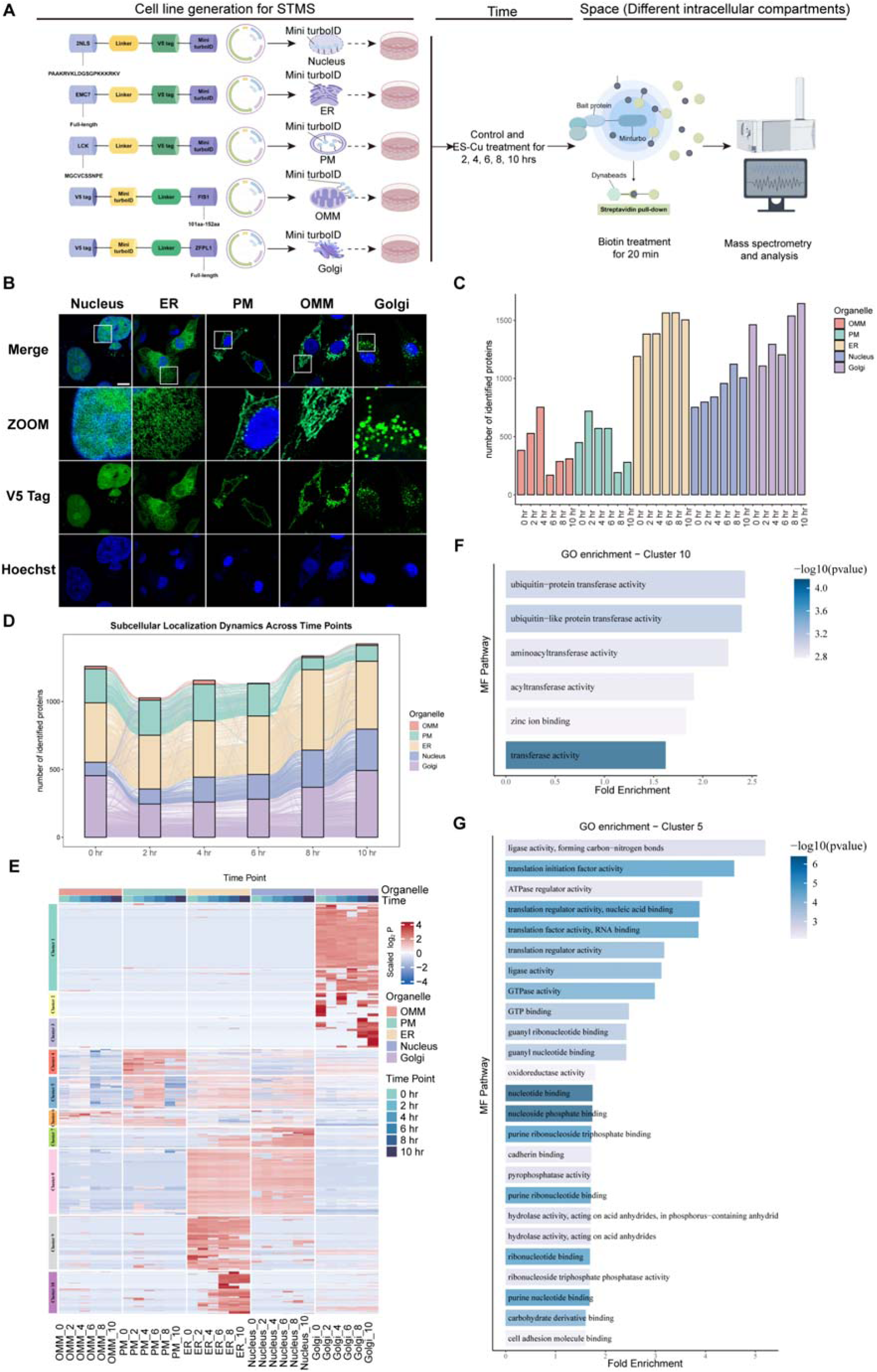
Spatio-Temporal Mass Spectrometry in living cells of early cuproptosis. (**A**) Schematic of Spatio-Temporal Mass Spectrometry in early cuproptosis. HT-1080 cells were treated with 1.5 μM CuSO_4_ and 15 nM ES at the indicated time points. (**B**) Expression and localization of miniTurboID in five organelles of HT-1080 cells. Scale bars, 10 µm. (**C**) The number of proteins in each organelle at different time points identified by mass spectrometry. (**D**) Sankey bar plot showing the changes of protein distribution among the organelles over time. The vertical bars represent the number of identified proteins in different organelles at various time points. Each color corresponds to a different organelle. The lines connecting the bars between different time points represent the transition of proteins between different organelles over time. The width of these lines indicates the quantity of proteins transition from one organelle to another. (**E**) Unsupervised clustering categorized subcellular proteome into 10 distinct clusters based on their dynamic expression pattern. (**F**) Enrichment analysis of genes in the tenth cluster using GO molecular function gene set with clusterProfiler. (**G**) Enrichment analysis of genes in the fifth cluster using GO molecular function gene set with clusterProfiler.

To profile subcellular proteome dynamics during early cuproptosis, we treated cells with copper ionophore Elesclomol and CuSO_4_ (ES-Cu), and collected protein samples at six time points for miniTurbo processing (Fig.1A). Biotinylated proteins were captured by streptavidin affinity purification, detected by streptavidin-HRP blotting (fig. S1A) and identified through mass spectrometry. Our spatial-temporal proteomic analysis identified over 4000 proteins across all organelles, with 2867 proteins retained after filtering for downstream analysis (table S1). Protein expression profiles demonstrated high consistency among replicates and revealed dynamic, organelle-specific expression patterns over time (fig. S1B and C).

We first analyzed the total number and abundance of proteins captured from each organelle at the six time points following ES-Cu treatment. While protein abundance in the ER was gradually decreased, other organelles exhibited no significant changes (fig. S1D). Notably, a sharp increase in protein numbers was observed at the PM and OMM within the first 2 hrs (Fig. 1C), followed by a steep decline to less than 50% of initial levels by 6 or 8 hrs post-treatment. In contrast, protein numbers in the ER and nucleus steadily increased, whereas the Golgi remained relatively stable (Fig. 1C). Analysis of differentially expressed proteins (DEPs) across organelles over time further revealed distinct organelle-specific patterns. The nucleus and ER showed a progressive increase in the number of upregulated proteins, whereas the OMM and PM exhibited a biphasic response characterized by an early surge in upregulated proteins followed by a later increase in downregulated proteins (fig S2, table S2).

We next examined changes of protein distribution among organelles following ES-Cu treatment. To define protein localization, we implemented a confidence scoring system (see Methods for details). At baseline (0 hr), protein localization based on our method was highly consistent with annotations in public databases (fig S2 F-H). Proteins with high confidence scores and compartment-specific localization were selected for subcellular trafficking analysis (table S3). While approximately 80% of proteins maintained their original localization over time, a subset exhibited dynamic relocalization, supporting the occurrence of protein trafficking during early cuproptosis (Fig. 1D).

To further characterize proteome dynamics, we categorized the subcellular proteome into 10 clusters based on their spatiotemporal expression patterns (Fig. 1E). Cluster 1, 2, 3, 9 and 10 exhibited organelle-specific localization, whereas other clusters included proteins distributed across multiple compartments. Most clusters displayed noticeable expression changes in response to ES-Cu treatment over time and were enriched with proteins associated with distinct molecular functions (Fig. 1, F and G and fig. S1E). For example, proteins in cluster 10 showed increasing expression in the ER from 4 hrs onward and were enriched for molecular functions related to ubiquitin-protein transferase activity, aminoacyltransferase activity, and zinc ion binding (Fig. 1F). These findings suggest that elevated copper levels induce proteotoxic stress in the ER, activate the ubiquitin-proteasome system and potentially interfere with zinc-dependent biological processes. Proteins in cluster 5 exhibited dynamic trafficking, with increased expression observed at the PM and OMM between 2 to 6 hrs, followed by a decline from 6 to 10 hrs. Simultaneously, their expression in the nucleus gradually increased, suggesting active inter-organelle protein redistribution (Fig. 1E). Function enrichment analysis of cluster 5 revealed associations with translation regulation, GTPase activity, nucleotide binding, and oxidoreductase activity (Fig. 1G). These results indicate that early copper-induced stress disrupts translation and promotes mitochondrial metabolic reprogramming. Proteins in cluster 7 were primarily localized to the ER and nucleus, with progressively increasing expression levels over time. These proteins were enriched for iron-sulfur (Fe-S) cluster binding, acetylation-dependent protein binding, and ubiquitin-protein transferase activity (fig. S1E). These findings suggest that copper stress destabilizes Fe-S cluster proteins, impairs mitochondrial function, and triggers acetylation-dependent protein regulation. The progressive nuclear accumulation of these proteins further supports a role for transcriptional adaptation in response to copper stress. Together, these findings highlight the mitochondria, ER and nucleus as central organelles involved in early response to copper-induced stress. Activation of the ubiquitin-proteasome system, destabilization of Fe-S cluster proteins, disruption of translation and dynamic protein trafficking represent key molecular events driving the initiation of copper-induced cell death.

### Loss of OXA1L in early cuproptosis impairs translation of oxidative phosphorylation proteins and activates UPR^mt^ through MRPL20 nuclear translocation

We hypothesize that proteins which show significant changes in the abundance and/or subcellular distribution may have functional impacts on cuproptosis. We selected 276 top-ranking DEPs (Fig 2A, fig S2) with additional 19 proteins potentially involved in cuproptosis based on published studies, resulting in a total of 295 candidate proteins as targets for a genetic functional screen to identify the molecular regulator of cuproptosis (table S4). There are 28 candidate genes whose depletion by shRNA and/or over-expression resulted in a change in cell viability following ES-Cu treatment by more than 30% (Fig. 2B and table S4).

**Fig. 2.**
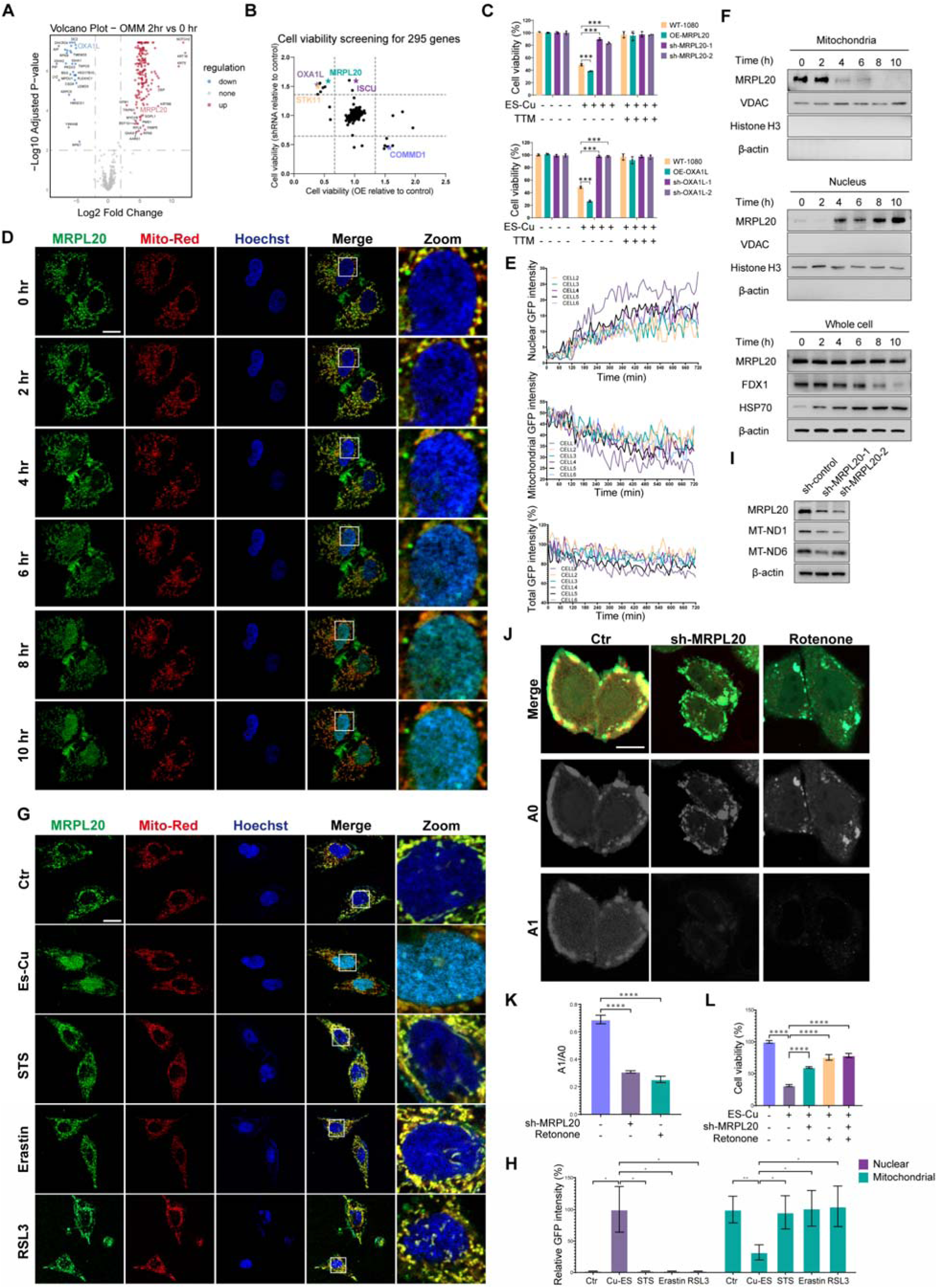
MRPL20 nuclear translocation occurs in early cuproptosis. (**A**) Volcano plot showing differentially expressed genes in the outer mitochondrial membrane (OMM) at 2-hr upon ES-Cu treatment compared to 0-hr. (**B**) A genetic functional screen identified the potential regulators of cuproptosis. HT-1080 cells were treated with 1.5 μM CuSO_4_ and 15 nM ES for 10 hrs. Genes whose overexpression (OE) or shRNA knockdown altered cell viability by more than 30% were selected for further validation. (**C**) Cell viability showing the effects of overexpression or knockdown of MRPL20 and OXA1L in HT-1080 cells on cuproptosis. Cells were treated as (B) in the presence or absence of 20 μM TTM. Data are presented as mean ± SD, n = 3. (**D**) Fluorescence images of mitochondrial and nuclear MRPL20-GFP expression in HT-1080 cells. Cells overexpressing GFP-tagged MRPL20 were treated with 1.5 μM CuSO_4_ and 15 nM ES and subject to image analysis at the indicated time points. Scale bars, 10 µm. (**E**) Histograms showing quantification of MRPL20-GFP intensity. (**F**)Western blot images showing endogenous MRPL20 expression in mitochondrial and nuclear fraction as well as the whole cell lysates. Cells were treated as (D). VDAC, Histone H3 and β-actin were used as the markers of mitochondria, nucleus and cytoplasm, respectively. (**G**) Fluorescence images of mitochondrial and nuclear MRPL20-GFP in HT-1080 cells upon treatment with cell death inducers. Cells overexpressing GFP-tagged MRPL20 were treated with cuproptosis inducer (1.5 μM CuSO_4_ and 15 nM ES for 10 hrs), apoptosis inducer (200 nM Staurosporine (STS), for 8 hrs), or ferroptosis inducers (50 μM Erastin for 24 hrs or 200 nM RSL3 for 8 hrs). Scale bars, 10 µm. (**H**) Histograms showing quantification of MRPL20-GFP intensity. Data are presented as mean ± SD, n = 4. (**I**) Western blot images showing the expression of MT-NDs in HT-1080 cells after knockdown of MRPL20 by shRNA. β-actin is used as a loading control. (**J**) Fluorescence image analysis of NAD+/NADH ratio in living cells. SoNar expressing HT-1080 cells were either transfected with MRPL20 shRNA or treated with 0.1 μM rotenone for 12 hrs. The images were pseudocolored with the fluorescence excited at 420 and 485 nm (A0-NADH, F485 nm; A1-NAD+, F420 nm). Scale bars, 10 µm. (**K**) Histogram showing the ratio of fluorescence intensities excited at A1 to A0. Data are presented as mean ± SD, n = 3. (**L**) Viability of HT-1080 cells with shRNA-mediated MRPL20 knockdown compared to control cells upon different treatments. Cells were treated with 1.5 μM CuSO_4_ and 15 nM ES in the presence or absence of 0.1 mM rotenone for 12 hrs. Data are presented as mean ± SD, n = 3. Statistical analysis was performed using a one-way ANOVA (C), (H), (K) and (L). * *p*-value < 0.05, ** *p*-value < 0.01, *** *p*-value < 0.001, **** *p*-value < 0.0001.

Two candidate genes, OXA1L and MRPL20, are involved in mitochondrial quality control^12^. The spatial-temporal proteomics analysis revealed a rapid decrease in mitochondrial OXA1L abundance, while MRPL20 exhibited a transient enrichment at the OMM before progressively accumulating in the nucleus over time (fig. S3A). We confirmed that knocking down OXA1L or MRPL20 prevented ES-Cu-induced cuproptosis whereas their overexpression increased cellular susceptibility to ES-Cu cytotoxicity (Fig. 2C). These effects were abolished by tetrathiomolybdate (TTM), a copper chelator in clinical trials for treatment of Wilson’s disease^13^, supporting a role for OXA1L and MRPL20 in mediating the cellular response to copper overload (Fig. 2C).

To confirm the nuclear translocation of MRPL20 during cuproptosis, we generated a HT1080 cell line overexpressing GFP-tagged MRPL20. The time-lapse confocal imaging showed an exclusive mitochondrial localization of MRPL20 before treatment (Fig. 2D and E). Upon ES-Cu exposure, MRPL20 signal intensity in the mitochondria rapidly declined accompanied with its nuclear accumulation. However, total MRPL20 abundance, as measured by total GFP intensity, exhibited only a modest reduction (Fig. 2D and E). Immunoblotting confirmed a time-dependent decrease of MRPL20 in the mitochondrial fraction and a concomitant increase in the nuclear fraction (Fig. 2F). This ES-Cu-induced mitochondria-nucleus trafficking can be accelerated by increasing ES concentration (fig. S3B and S3C). Immunofluorescent staining of endogenous MRPL20 corroborates the finding in MRPL20-overexpressing cells (fig. S3D and S3E). Moreover, MRPL20 nuclear translocation was blocked by TTM (fig. S3D and S3E), firmly supporting that this response is associated with copper influx.

To confirm this nuclear translocation is a specific response to copper stress, HT-1080 cells were treated with apoptosis inducer (Staurosporine, STS)^14^ or ferroptosis inducers (Erastin and RSL3)^15^ (Fig. 2G and 2H). Indeed, nuclear accumulation of MRPL20 occurred exclusively in the response to cuproptosis inducer (Fig. 2G and 2H), confirming its specificity to copper-induced cell death.

Given its role as a mitochondrial ribosomal protein^16^, we speculate that MRPL20 deficiency may impair mitochondrial oxidative phosphorylation (OXPHOS) protein synthesis. Genes encoding the core subunits of OXPHOS complex I, including mitochondrially encoded NADH dehydrogenase 1-6 (Mt-DN1-6) are located in mitochondrial DNA and rely on mitochondrial ribosomes for translation^17^. MRPL20 knockdown decreased Mt-ND protein abundance (Fig. 2I). Moreover, the intracellular ratio of NAD+/NADH was decreased upon MRPL20 depletion, comparable to the effect of Rotenone, an inhibitor of ETC complex I (Fig. 2J and 2K). Similar to MRPL20 knockdown, inhibition of complex I activity by rotenone also mitigated ES-Cu cytotoxicity (Fig. 2L), underscoring the requirement of mitochondrial OXPHOS activity for cuproptosis induction. Notably, the combination of rotenone and MRPL20 knockdown did not further improve cell viability, suggesting that their actions either overlap or share a common pathway (Fig. 2L). Collectively, we demonstrate that mitochondrial MRPL20 is essential for mRNA translation of the mitochondrial electron transport chain complex I, and mitochondrial respiratory activity.

OXA1L plays a crucial role in inserting mitochondrial proteins encoded by either mitochondrial or nuclear genome, into the inner mitochondrial membrane (IMM)^18^. Its C-terminal matrix-exposed domain interacts with the large mitochondrial ribosome subunit proteins including MRPL20^12^, to facilitate the translocation of newly synthesized OXPHOS proteins to the IMM^19^. Spatial-temporal proteomic analysis identified a reduction in mitochondrial OXA1L upon ES-Cu treatment (Fig. 2A and fig. S3A), a finding confirmed by the immunoblotting (Fig. 3A). Loss of OXA1L has been reported to reduce mitochondrial OXPHOS complexes^20^. Consistently, depletion of OXA1L by shRNA or ES-Cu treatment decreased the abundance of OXPHOS complex I (Fig. 3B and 3C).

**Fig. 3.**
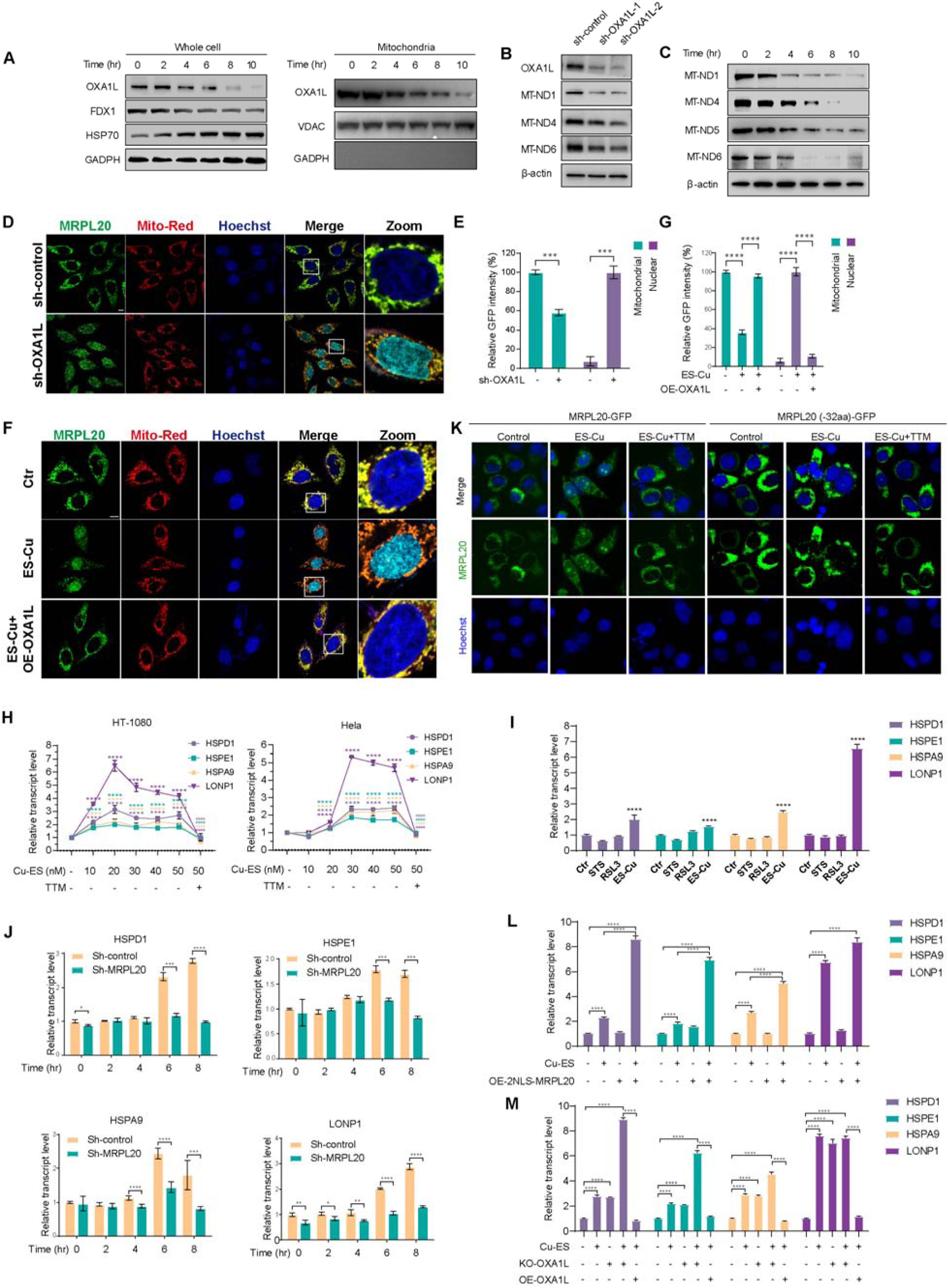
Loss of OXA1L in early cuproptosis promotes MRPL20 mitochondrial-nuclear trafficking and activates UPR^mt^. (**A**) Western blot images showing the expression of OXA1L in whole cell lysates and mitochondrial fraction of HT-1080 cells. Cells were treated with 1.5 μM CuSO_4_ and 15 nM ES for different time periods as indicated. GAPDH and VDAC were used as the markers of cytoplasm and mitochondria, respectively. (**B** and **C**) Western blot images showing the expression of MT-NDs in HT-1080 cells with shRNA-mediated OXA1L knockdown (B) or upon ES-Cu treatment (C). In (C), cells were treated as (A). β-actin is used as a loading control. (**D** and **E**) Fluorescence images of mitochondrial and nuclear MRPL20-GFP expression in HT-1080 cells with OXA1L knockdown. The quantification of MRPL20-GFP intensity was shown in (E). (**F** and **G**) Fluorescence images of mitochondrial and nuclear MRPL20-GFP expression in HT-1080 cells. Cells were treated with 1.5 μM CuSO_4_ and 15 nM ES in the presence or absence of OXA1L overexpression. The quantification of MRPL20-GFP intensity was shown in (G). Scale bars, 10 µm. Data are presented as mean ± SEM, n = 3 (D), n = 4 (E). (**H**) Relative transcriptional levels of UPR^mt^ genes in Hela and HT-1080 cells after treatment with 1.5 μM CuSO_4_ and different concentrations of ES for 12 hrs measured by qPCR. Data are presented as mean ± SEM, n = 4. (**I**) Relative transcriptional levels of UPR^mt^ genes in HT-1080 cells after treatment with 1.5 μM CuSO_4_ and 15 nM ES, 200 nM STS or 200 nM RSL3 for 8 hrs. Data are presented as mean ± SEM, n = 4. (**J**) Relative transcriptional levels of UPR^mt^ genes in HT-1080 cells transfected with MRPL20 shRNA or control shRNA followed by treatment with 1.5 μM CuSO_4_ and 20 nM ES for different time periods as indicated. Data are presented as mean ± SEM, n = 3. (**K**) Fluorescence images of mitochondrial and nuclear MRPL20-GFP expression in HT-1080 cells overexpressing GFP-tagged MRPL20 in full length or with NLS deletion upon treatment with1.5 μM CuSO_4_ and 15 nM ES in the presence or absence of 20 μM TTM for 12 hrs. (**L**) Relative transcriptional levels of UPR^mt^ genes in 2NLS-MRPL20 overexpressing HT-1080 cells after treatment with or without 1.5 μM CuSO_4_ and 20 nM ES for 8 hrs. Data are presented as mean ± SEM, n = 4. (**M**) Relative transcriptional levels of UPR^mt^ genes in HT-1080 cells under different conditions. The treatment groups include: (1) Control (No treatment), (2) Cu-ES treatment, (3) OXA1L KO, (4) OXA1L KO cells treated with Cu-ES, and (5) OXA1L overexpressing cells treated with Cu-ES. Data are presented as mean ± SEM, n = 4. Statistical analysis was performed using a one-way ANOVA (H), (I), (J), (L) and (M) and two-tailed unpaired Student’s t test (E) and (G). * *p*-value < 0.05, ** *p*-value < 0.01, *** *p*-value < 0.001, **** *p*-value < 0.0001.

Disrupting OXPHOS can impair mitochondrial protein import efficiency and activate the retrograde mitochondrial-nucleus communication, triggering mitochondrial unfolded protein response (UPR^mt^), a nuclear transcriptional response that induce expression of mitochondrial chaperones and proteases^21^. We thus hypothesize that loss of OXA1L induces UPR^mt^ by promoting MRPL20 nuclear translocation. As expected, OXA1L knockdown reduced mitochondrial MRPL20 level and promoted its nuclear accumulation (Fig. 3D and 3E). Conversely, over-expression of OXA1L retained MRPL20 in mitochondria even in the presence of ES-Cu (Fig. 3F and 3G), demonstrating that OXA1L is critically required for MRPL20 mitochondrial localization and its function in OXPHOS protein synthesis.

To determine UPR^mt^ activation under copper stress, we firstly examined the mRNA expression levels of four genes (HSPD1, HSPE1, HSPA9 and LONP1), previously reported to be induced during UPR^mt22^. Indeed, ES-Cu treatment led to a dose-dependent increase in their expression, which was effectively blocked by TTM (Fig. 3H). Moreover, transcriptional activation of these genes was only be induced by ES-Cu not by apoptosis (STS) or ferroptosis (RSL3) inducers (Fig. 3I), confirming that UPR^mt^ is a specific response to copper stress. Importantly, MRPL20 knockdown impaired ES-Cu-induced UPR^mt^ activation, supporting its critical role in this response (Fig. 3J).

The ability of MRPL20 to translocate to the nucleus under copper stress suggests the presence of an intrinsic nuclear-localization signal (NLS). Using the cNLS Mapper tool^23^, we identified three potential NLS motifs (fig. S3F). Deletion of one motif (RHFRGRKNRCYRLAVRTVIRAFVKCTKARYLK) completely abolished MRPL20 nuclear translocation upon ES-Cu treatment (Fig. 3K), highlighting its essential role in cooper-induced nuclear import.

To further investigate the impact of MRPL20 nuclear translocation, we generated a construct with an exogenous NLS fused to MRPL20, forcing its nuclear translocation under normal condition (fig. S3G). In the absence of copper stress, forced nuclear translocation of MRPL20 alone did not activate UPR^mt^ gene transcription, but significantly enhanced the transcriptional response upon ES-Cu treatment (Fig. 3L). These findings suggest that MRPL20 nuclear trafficking is necessary but not sufficient to activate UPR^mt^.

By contrast, OXA1L depletion alone activated UPR^mt^ gene transcription to at a level comparable to ES-Cu treatment, while overexpression of OXA1L completely abolished UPR^mt^ activation (Fig. 3M). Together, our findings revealed a previously unrecognized role of OXA1L in mediating copper stress-induced-UPR^mt^ activation.

### AMPK-mediated ISCU phosphorylation under copper stress promotes cuproptosis

Another interesting candidate gene identified in the genetic functional screen is STK11, whose overexpression increased ES-Cu sensitivity, whereas depletion by shRNA had the opposite effect (Fig. 2B and table S4). It encodes the protein LKB1, a key kinase that phosphorylates and activates AMPK, a central energy sensor^24^. This finding suggests that defects in mitochondrial OXPHOS during early cuproptosis may trigger AMPK activation. Consistent with this hypothesis, immunoblotting revealed a time-dependent increase in AMPK phosphorylation at threonine 172 following ES-Cu treatment (fig. S4A). Similarly, AMPK activation via glycolysis inhibition with 2-Deoxy-D-glucose (2DG)^25^, increased sensitivity to ES-Cu treatment and promoted cuproptosis (Fig. 4A). Conversely, inhibition of AMPK activity by AMPK-IN-3^26^ rendered cells resistant to ES-Cu-induced cytotoxicity (Fig. 4A). These results demonstrate that AMPK signaling is required for cuproptosis induction.

**Fig. 4.**
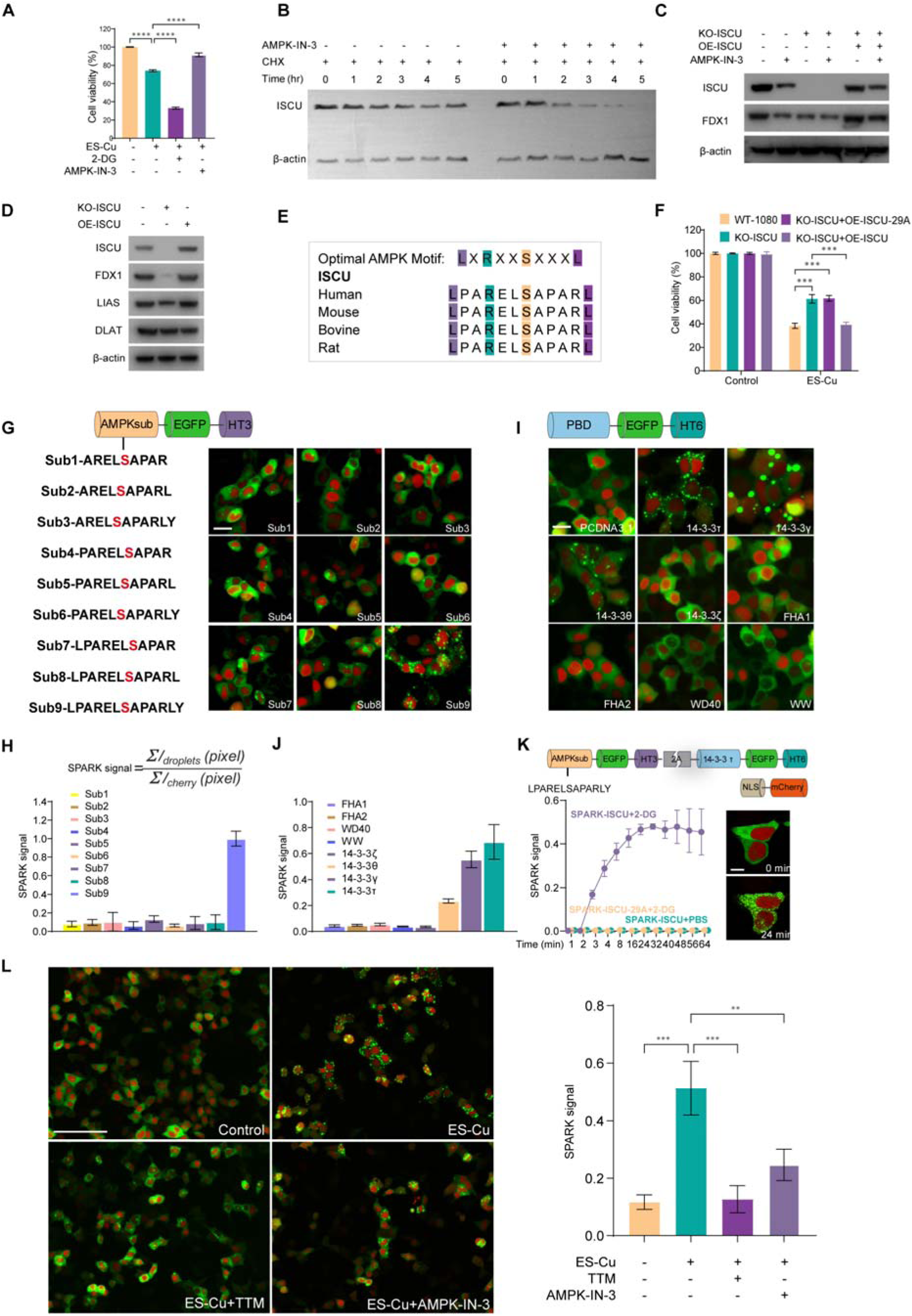
AMPK-mediated ISCU phosphorylation under copper stress promotes cuproptosis. (**A**) Viability of HT-1080 cells upon different treatments. Cells were treated with 1.5 μM CuSO_4_ and 15 nM ES alone or in combination with 50 μM 2-Deoxy-D-glucose (2-DG) or 25 μM AMPK-IN-3 for 24 hrs. Data are presented as mean ± SD, n = 3. (**B**) Western blot analysis of ISCU protein levels in HT-1080 cells pre-treated with 50 μM cycloheximide (CHX) for 30 minutes, followed by treatment with 25 μM AMPK-IN-3 treatment for the indicated durations. (**C**) Western blot analysis of the expression of Fe-S cluster-containing proteins in in HT-1080 cells with ISCU knockout or overexpression in the presence or absence of 25 μM AMPK-IN-3 for 12 hrs. (**D**) Western blot analysis of the expression of Fe-S cluster-containing proteins in in HT-1080 cells. (**E**) Amino acid sequence alignment of an AMPK consensus phosphorylation site motif of ISCU across species. (**F**) Viability of HT-1080 cells with ISCU knockout plus overexpression of ISCU wild type or S29A mutant upon Cu-ES treatment. Cells were treated with 1.5 μM CuSO_4_ and 15 nM ES for 24 hrs. Data are presented as mean ± SD, n = 3. (**G**) Design and selection of AMPK substrate peptides based on the AMPK consensus motif surrounding serine29 of human ISCU protein. HEK293T cells were transiently co-transfected with AMPK-SPARK plasmid plus NLS-mCherry plasmid and then treated with 50 μM 2-DG for 30 min. Scale bars, 20 µm. (**H**) The histogram shows GFP/mCherry intensity ratio of different AMPK substrate peptides. Data are presented as mean ± SD, n = 3. (**I**) Design and selection of the phosphoserine binding domain peptides. Cells were treated as (G). Scale bars, 20 µm. (**J**)The histogram showing GFP/mCherry intensity ratio of different phosphoserine binding domain peptides. Data are presented as mean ± SD, n = 3. (**K**) Schematic diagram of the optimized structure of AMPK-SPARK. Fluorescence images of HT-1080 cells expressing AMPK-SPARK containing S29 ISCU peptide before and after the addition of 50 μM 2-DG. The histogram showing quantification of AMPK-SPARK intensity. Data are presented as mean ± SD, n = 3. (**L**) Fluorescence images showing changes of AMPK-SPARK signal intensity in HEK293T upon different treatments. Cells were exposed to 1.5 μM CuSO_4_ and 15 nM ES with or without 20 μM TTM or 25 μM AMPK-IN-3 for 24 hrs. The chart aligned with fluorescent images. Scale bars, 125 µm. Data are presented as mean ± SD, n = 3. Statistical analysis was performed using a one-way ANOVA (A), (F) and (L). * *p*-value < 0.05, ** *p*-value < 0.01, *** *p*-value < 0.001, **** *p*-value < 0.0001.

Our STMS analysis identified alterations in proteins associated with Fe-S cluster binding (Fig. 1G). AMPK has been reported to phosphorylate Fe-S cluster assembly enzyme (ISCU), a scaffold protein required for Fe-S cluster biogenesis, at serine residues 14 and 29^27^. This phosphorylation is thought to promote ISCU binding to 14-3-3 proteins, thereby stabilizing ISCU protein and enhancing Fe-S cluster synthesis, which in turn maintains redox and energy homeostasis^27^. We found that AMPK inhibition, indeed, markedly reduced ISCU level in the presence of cycloheximide (Fig. 4B), highlighting the role of AMPK signaling activity in stabilizing ISCU. We further examined the levels of Fe-S cluster-containing proteins, including FDX1 and LIAS, in cells with ISCU knockout or treated with AMPK-IN-3. As expected, loss of ISCU or AMPK inhibition diminished FDX1 and LIAS protein expression levels (Fig 4C and 4D).

Sequence analysis confirmed that the serine 29 (S29) of ISCU is situated within a highly conserved AMPK consensus motif across species, supporting its role as an AMPK specific phosphorylation site (Fig. 4E). Notably, the genetic functional screen showed that ISCU depletion by shRNA reduced ES-Cu sensitivity, consistent with the effects of STK11 knockdown or AMPK inhibition (Fig. 2B and 4A and table S4). We thus proposed that ISCU, as an AMPK substrate, mediates AMPK signaling in early cuproptosis. To test this hypothesis, we overexpressed an ISCU non-phosphorylatable mutant by substituting the serine 29 residue with alanine in ISCU-depleted cells. Overexpression of this phosphorylation mutant failed to restore sensitivity to ES-Cu in ISCU-knockdown cells (Fig. 4F). Together, these findings suggest that AMPK-ISCU signaling enhances susceptibility to cuproptosis.

To monitor AMPK activation during cuproptosis, we employed SPARK (separation of phase-based activity reporter of kinase), a technology we developed for real-time visualization of PKA and ERK kinase activity in live cells^28^ (fig. S4B). To rationally design a reporter that can sense the phosphorylation status of ISCU at S29 by AMPK, we designed and screened nine AMPK-derived substrate peptides based on the ISCU S29 consensus motif. A peptide (LPARELSAPARLY) was selected and fused with EGFP (Fig. 4G and 4H), along with a trimeric homo-oligomeric tag (HOTag3). Subsequently, through domain optimization for binding, we selected 14-3-3 tau (Fig. 4I and 4J), which is a phosphoserine binding domain (PBD) and fused with hexameric homo-oligomeric tag (HOTag6), for AMPK activity-dependent protein-protein interaction. Finally, we employed a self-cleaving 2A sequence to construct both components together (Fig. 4K).

The intensely green liquid droplets were rapidly formed upon the addition of 2DG or starvation (fig. S4C and D) and this formation was blocked by AMPK-IN-3 (fig. S4E), indicating activation of AMPK-ISCU pathway in response to energy stress. Conversely, the ISCU S29A-mutated sensor failed to form the droplets upon 2DG treatment (Fig. 4K and fig. S4C and D), confirming the specificity of the ISCU S29 sensor. Since this AMPK-SPARK achieves a large dynamic range, high brightness, and fast kinetics for reporting AMPK activity, we exploited it to monitor AMPK activity in early cuproptosis. ES-Cu treatment led to formation of EGFP droplets, which was blocked by the copper chelator TTM and AMPK inhibitor AMPK-IN-3 (Fig. 4L). These results strongly support that AMPK signaling is activated in response to cupper stress.

### COMMD4 is a biomarker of cuproptosis

Our gene functional screen identified copper metabolism domain containing 1 (COMMD1) as a regulator of cuproptosis, with its overexpression promoting cell survival, whereas COMMD1 depletion sensitized cells to ES-Cu-induced cytotoxicity (Fig. 2B and table S4). Furthermore, STMS analysis revealed upregulation of COMMD4 upon ES-Cu treatment (fig. S5A), a finding confirmed by immunoblotting in the whole cell lysate (Fig. 5A). Both COMMD1 and COMMD4 are members of the COMMD family, which consists of ten evolutionarily conserved proteins characterized by a highly conserved carboxy-terminal COMM domain (fig. S5B). To systematically characterize their functional roles in cuproptosis, we generated a panel of HT1080 cell lines expressing COMMD1 to 10 fused with GFP (fig. S5B). Except COMMD1, overexpression of other COMMD family members did not improve cell viability under copper overload (Fig. 5B). Corroborating its role in facilitating copper excretion^29^, our findings reinforce COMMD1 as an important regulator of cuproptosis.

**Fig. 5.**
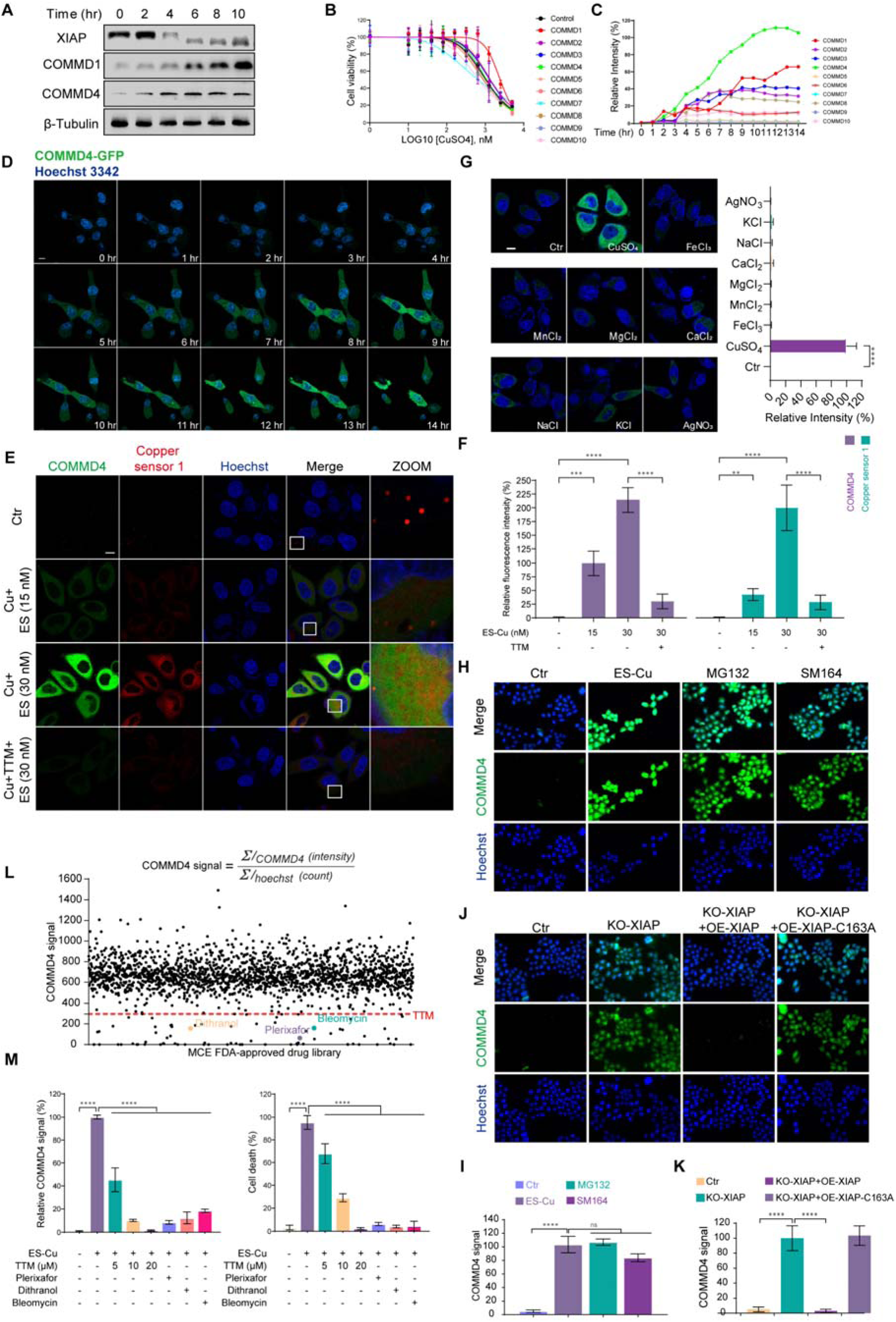
COMMD4 is a sensitive sensor for intracellular copper accumulation. (**A**) Western blot analysis of endogenous XIAP, COMMD1 and COMMD4 expression in HT-1080 cells treated with 1.5 μM CuSO_4_ and 15 nM ES for the different time periods as indicated. β-Tubulin is used as a loading control. (**B**) Cell viability analysis in HT-1080 cells transiently transfected with GFP-tagged COMMD family members upon treatment with different concentrations of CuSO_4_ and 15 nM ES for 24 hrs. Data are presented as mean ± SD, n = 4. (**C**) Quantification of GFP fluorescence intensity in HT-1080 cells transiently transfected with GFP-tagged COMMD family members upon treatment with 1.5 μM CuSO_4_ and 15 nM ES for different time periods as indicated. (**D**) Time lapses confocal images showing COMMD4-GFP expression upon treatment with 1.5 μM CuSO_4_ and 15 nM ES. Scale bars, 10 µm. (**E** and **F**) Fluorescence images of COMMD4-GFP and copper sensor 1 in HT-1080 cells treated with 1.5 μM CuSO_4_ and 15 or 30 nM ES in the presence or absence of 20 μM TTM for 12 hrs. Scale bars, 10 µm. The quantification of fluorescence intensity was shown in F. Data are presented as mean ± SD, n = 3. (**G**) Fluorescence images of COMMD4-GFP expression in HT-1080 cells upon treatment with CuSO_4_, FeCl3, MnCl2, MgCl2, NaCl and KCl at 250 μM, or CaCl2 and AgNO3 at 50 μM for 12 hrs. Scale bars, 10 µm. (**H** and **I**) Fluorescence images of COMMD4-GFP in HT-1080 cells treated with either 1.5 μM CuSO_4_ and 15 nM ES, 20 µM MG132 for 12 hrs or 10 nM SM164 for 24 hrs. Scale bars, 10 µm. The quantification of GFP signal intensity was shown in (I). (**J** and **K**) Fluorescence images of COMMD4-GFP in HT-1080 cells with XIAP knockout. The quantification of GFP signal intensity was shown in (K). (**L**) Quantification of COMMD4-GFP fluorescence intensity normalized to cell count in HeLa cells transiently transfected with COMMD4-GFP after treatment with 5 μM FDA approved drugs library in the presence of ES-Cu for 24 hrs. (**M**) Relative COMMD4-GFP fluorescence intensity and cell death in Hela cells overexpressing COMMD4-GFP upon treatment with Plerixafor, Dithranol, and Bleomycin at 5 μM in combination with 1.5 μM CuSO_4_ and 35 nM ES for 24 hrs compared to 5 μM (low dose, l), 10 μM (middle dose, m) and 20 μM (high dose, h) TTM. Data are presented as mean ± SD, n = 3. Statistical analysis was performed using a one-way ANOVA (F), (G), (M), (I) and (K). * *p*-value < 0.05, ** *p*-value < 0.01, *** *p*-value < 0.001, **** *p*-value < 0.0001.

On the other hand, COMMD4 displayed a distinct response to ES-Cu treatment, with a rapid, time-dependent increase in cytoplasmic signal intensity, significantly faster than other COMMD family members (Fig. 5C and 5D, fig. S5C). This Cu-induced COMMD4 upregulation was blocked by the copper chelator TTM (Fig. 5E and 5F) and was not observed with other metal ions (Fig. 5G). Strikingly, under normal condition, COMMD4-GFP overexpressing cells did not exhibit green fluorescence (Fig. 5D). Treatment with a proteasome inhibitor MG132, but not an autophagy inhibitor CQ, effectively restored COMMD4 expression, suggesting that COMMD4 is degraded via ubiquitination-proteasome pathway under basal condition (Fig. 5H and 5I, fig. S5D).

The conserved COMM domain serves as protein-protein interaction platform among COMMD family members and their binding partners. Previous studies have shown that XIAP mediates K48-linked polyubiquitination of COMMD1, targeting it for proteasome degradation. Copper overload promotes XIAP degradation, leading to increased COMMD1 stability^30^. We speculate that a similar mechanism accounting for COMMD4 degradation. Immunoblotting confirmed copper stress led to decrease of XIAP expression accompanied with increase of COMMD1 and COMMD4 expression (Fig. 5A). XIAP knockout or inhibition by SM164, a Smac mimetic^31^,restored COMMD4 expression, an effect which can be blocked by overexpressing XIAP, but not its C163A mutant, in KO cells (Fig. 5H - 5K). We thus provide the direct evidence supporting XIAP-mediated COMMD4 degradation under basal condition.

Copper sensor-1 (CS1) is commonly used fluorescent dye for imaging labile copper in living cells^28, 32^. Despite its high sensitivity, it has several limitations, including photobleaching, limited stability, fluorescence quenching and high cost. Compared to CS1, COMMD4-GFP exhibited similar sensitivity to intracellular copper fluctuation (Fig. 5E and 5F). Notably, COMMD4 displayed a stronger signal than CS1 at a low dose of ES (15 nM), suggesting greater sensitivity to copper flux (Fig. 5F). Given its low background signal, high sensitivity and non-cytotoxicity natures, COMMD4 represents a promising biomarker for cuproptosis detection.

We thus determined to use this GFP-tagged COMMD4 cell model for a high-content drug screening for new cuproptosis inhibitors. Cells were treated with 2,001 FDA-approved drugs in the presence of ES-Cu. The copper stress was assessed by GFP fluorescence intensity and cell death was detected by propidium iodide (PI) staining (fig. S5E). TTM, as a copper chelating agent currently in Phase II clinical trials for oncology^33^, served as a positive control.

Three therapeutic agents, namely Plerixafor, Dithranol and Bleomycin, exhibited a significantly stronger protective effect against ES-Cu-induced cytotoxicity than TTM (Fig. 5L and 5M, fig. S5F). Importantly, these compounds specifically inhibited cuproptosis, as they did not affect apoptosis (STS-induced) or ferroptosis (Erastin and RSL3-induced) (fig. S5G). Thus, using COMMD4 as a unique sensor of cuproptosis, we successfully identified new cuproptosis inhibitors.

## Discussion

Cuproptosis is a copper-dependent mitochondrial cell death driven by proteotoxic stress due to aggregation of lipoylated TCA cycle proteins and degradation of Fe-S cluster proteins^4^. Although important advances have been made in understanding the roles of copper in human diseases and mechanisms of cuproptosis ^34–38^, the questions that what the early effectors of cuproptosis are and how cells defense against copper-induced damages are still largely unanswered. Furthermore, the lack of specific and reliable biomarkers for cuproptosis poses a major limitation in the field. To address these questions, we conducted Spatio-Temporal Mass Spectrometry (STMS) through a series of miniTurboID-based proximity labelling over time followed by the quantitative proteomic analysis. We demonstrate that cuproptosis is a mitochondria-centered process that impacts multiple organelles through the regulation of protein abundance and trafficking. Subsequently through a genetic functional screen and target validation based on genetic methods and confocal live cell imaging, we discover the molecules and signalling pathways that are key regulators of cuproptosis.

We identify loss of OXA1L promoting translocation of MRPL20 from mitochondria to nuclei as an early adaptive response to copper stress. Given the critical role of OXA1L in translation, insertion and assembly of OXPHOS proteins^18^, loss of OXA1L perturbates MRPL20 importing into mitochondria, which markedly impairs complex I formation and function. This copper-overload-induced mitochondrial dysfunction activates mitochondrial unfolded protein response (UPR^mt^), a nuclear transcriptional response to induce expression of genes to re-establish mitochondrial proteostasis and function^22^. Our findings thus define OXA1L as the key regulator of cuproptosis-related UPR^mt^ and OXA1L-MRPL20 as a critical signaling event involved in retrograde mitochondria-to-nucleus communication and thereby protect from cupper-induced proteotoxic stress (Fig. 6).

**Fig. 6.**
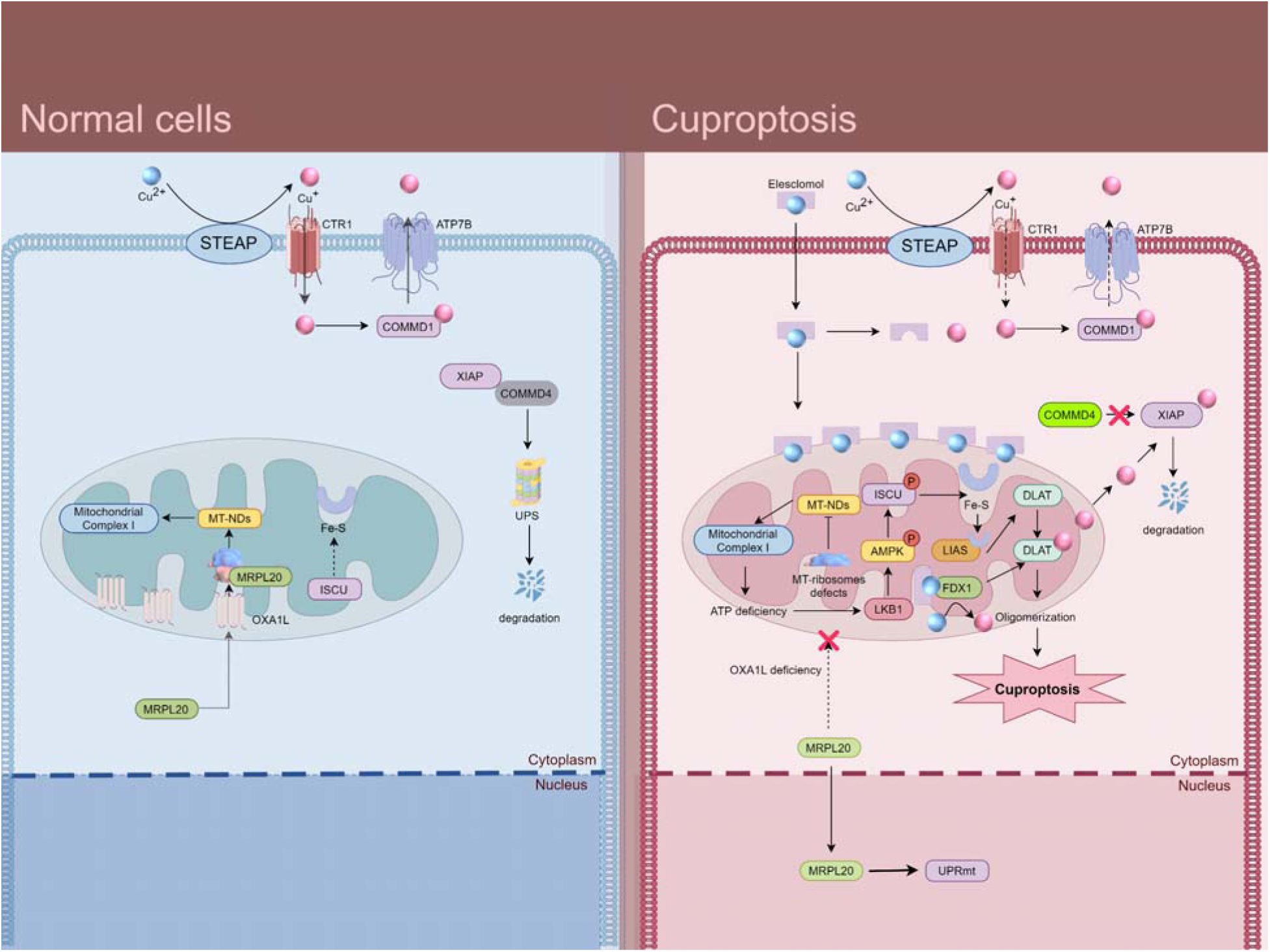
Schematic of new mechanisms of cuproptosis.

We also identified early activation of LKB1-AMPK-ISCU signaling increases stability of Fe-S containing proteins but enhances cells susceptibility to cuproptosis. Mitochondrial proteotoxic stress impairs TCA and OXPHOS activities and consequently reduces ATP production. This energy stress activates LKB1-AMPK signaling. AMPK subsequently phosphorylates ISCU at S29 which promotes ISCU binding to 14-3-3s. This binding stabilizes ISCU protein and enhances synthesis of Fe-S clusters and consequently the abundance of Fe-S-containing mitochondrial proteins, including FDX1 and LIAS (Fig. 6). LIAS catalyzes the formation of lipolated dihydrolipoamide S-acetyltransferase (DLAT) whose aggregation causes mitochondrial proteotoxic stress and cuproptosis (Fig. 6). These findings are in line with the role of AMPK signalling in protection of mitochondrial proteostasis by inhibiting misfolded cytosolic proteins importing into mitochondria recently identified in yeast^21^. Collectively, we provide evidence for AMPK as a critical metabolic checkpoint, regulating mitochondrial proteotoxic stress during cuproptosis. Based on these mechanistic discoveries, we developed a robust AMPK-SPARK, an EGFP phase separation-based activity reporter of AMPK for imaging endogenous AMPK activity in living cells during cuproptosis. Because of its large dynamic range and bright fluorescence, this system can be used to high content screening for genes or chemicals that affect AMPK activity. Furthermore, the reversibility of SPARK signal and no apparent toxicity in transgenic animals without perturbing animal development^28^, enable analysis of AMPK signaling and assessment of cuproptosis in a quantitative way in vivo.

A notable limitation in research of cuproptosis is the lack of specific and reliable biomarkers for early detecting intracellular copper deregulation. We develop a COMMD4-GFP reporter to monitor cuproptosis in living cells. Under normal condition, COMMD4 is rapidly degraded via XIAP-mediated ubiquitination, resulting in its signal undetectable. Upon copper flux, reduced XIAP led to accumulation of COMMD4, resulting in a significant increase of fluorescence intensity. This unique characteristic offers a wide dynamic range of signal detection. Together with its specificity, sensitivity and non-cytotoxicity, making it a superior and more practical approach for drug screening applications. Using COMMD4 as a sensor of cuproptosis, we identified three pharmacological agents (Plerixafor, Dithranol, and Bleomycin) as potential cuproptosis inhibitors with greater potency than TTM. Our findings thus may pave the new way for drug repurposing in the treatment of diseases related to copper disorders, such as Wilson’s and Menkes diseases, as well as cancer.

Taken together, this study delineates the spatiotemporal subcellular protein profiles in early cuproptosis, deciphers the molecular mechanisms underlying cellular adaptive response against copper-induced mitochondrial proteotoxic stress, including OXA1L-MRPL20-driven mitochondria-nucleus communication and AMPK-ISCU signaling-mediated Fe-S cluster synthesis and establishes COMMD4 as a specific sensor of cuproptosis. Our work not only expand current understanding of cuproptosis but also provide molecular tools and data resource for further research in this area.

## Methods

### Plasmids

All oligonucleotides (table S5) were synthetized by Beijing tsingke Biotech Co., Ltd. (Beijing, China). Plasmid pcDNA3.1(+), pCDH-CMV-EF1α, pLKO.1 and lentiCRISPR v2 were purchased from Addgene. SoNar plasmid was likewise synthetized by Beijing tsingke Biotech.

To engineer the plasmids for overexpression of genes in the functional screen, full-length cDNA sequences of the candidate genes were amplified from HT-1080 cells cDNA library via PCR and ligated into the pcDNA3.1(+) or pCDH-CMV-EF1α which contained EGFP and featured NheI and ECORI restriction sites using one step cloning kit (C112, Vazyme).

To generate the shRNA plasmids, the shRNA oligonucleotides were firstly annealed by PCR. The resulting double-stranded oligonucleotides were then ligated into the pLKO.1 vector using AgeI and ECORI restriction sites with T4 DNA ligase (C301, Vazyme).

To construct the CRISPR knockout plasmids, annealed sgRNA oligonucleotides were ligated into the lentiCRISPR v2 vector using BsmBI-v2 restriction sites by T4 DNA ligase.

To build miniTurboID plasmid, V5-tag and miniTurboID were tandemly fused with different organelle-specific bait protein markers or signal sequences and ligated into pCDH-CMV-EF1α.

To build AMPK-SPARK plasmid, AMPK substrate peptides and phosphoserine binding domain were connected by trimeric homo-oligomeric tag (HT3), self-cleaving 2A sequence and hexameric homo-oligomeric tag (HT6) and ligated into pcDNA3.1(+).

### Cell Culture and Generation of Stable Cell Lines

HT-1080, HEK293T, A549 and Hela cells purchased sourced from Procell Life Science & Technology were cultured in DMEM supplemented with 10% fetal bovine serum (FCS500, ExCell) at 37 in 5% CO_2_. They were confirmed to be mycoplasma negative.

For transient overexpression or knockdown, calcium phosphate transfection reagent (21 mM HEPES, 137 mM NaCl, 5.55 µM Dextrose, 50 mM KCl, 70 mM Na_2_HPO_4_, 2.5 M CaCl_2_, PH 7.1) or Hieff Trans (40802ES03, YEASEN) were used according to the manufacturer’s protocol.

To generate stable cell lines, two lentiviral packaging vectors (VSVG and psPAX2) and the TuroID expression plasmids were co-transfected into HEK293T cells. Lentiviral supernatants were collected at 48 hrs post-transfection and infected HT-1080 cells in the presence of 4 µg/mL polybrene. Cells were selected by 0.2–1 µg/mL puromycin or 10 µg/mL blasticidin depending on the expression plasmids transduced, for 1 week.

### Cell viability assay

Cells were seeded onto 96-well plates at a density of 1 × 10^4^ cells in 100 μL medium per well. After overnight incubation, 100 μL drug dilutions were added to the cell culture. Following the designated treatment period, 10 μL of CCK-8 reagent (K1018, APExBio) was added into each well containing 200 μL of medium. After incubation for 1 hr, the absorbance was measured at 450 nm using a Multiskan SkyHigh Microplate Spectrophotometer (Thermo Fisher Scientific). Cell viability was calculated by comparing the absorbance of cells treated with ES-Cu to that of untreated cells. In the gene functional screen, genes whose overexpression or knockdown altered cell viability by more than 30% were selected as the potential regulator of cuproptosis for further validation.

### High content screening for FDA-approved drug library

Hela cells overexpressing COMMD4-GFP were seeded onto 96-well plates at a density of 1× 10^4^ cells in 100 μL per well. After overnight incubation, cells were exposed to 5 μM FDA-approved drugs library (HY-L022, MCE) in combination with 2.5 μM CuSO_4_ and 35 nM ES for 24 hrs. Then 10 μg/mL Hoechst 33342 and 50 μg/mL PI were added into the medium and incubated for 30 min. Images were acquired by EVOS™ M7000 Imaging Systems (Thermo Fisher Scientific) and analyzed using Celleste Image Analysis Software (Thermo Fisher Scientific). Intracellular copper level was measured by the GFP fluorescence intensity/ cell count. Cell death was measured by the PI + cell count/ total cell count.

### Live cell imaging

Cells were plated onto a 35 mm confocal dish (J40204, Jingan) in DMEM with 10% FBS until reaching 50% confluence. After treatment with Cu-ES, the time-lapse images were obtained from a laser-scanning confocal microscope (LSM Zeiss 900, Zeiss) inverted confocal laser scanner microscope with 405, 488, 561, or 640 nm wavelength lasers.

### Immunofluorescence

Cells were plated onto 35 mm confocal dishes. After incubation overnight, cells were fixed with 4% paraformaldehyde and gently agitated on a low-speed shaker for 15-20 min. Fixed cells were permeabilized with 5% (v/v) Triton X-100 for 10 min, washed three times with PBS for 5 min each and blocked with 2% BSA at RT for 30 min. Cells were incubated with anti-MRPL20 antibody (1:300, DF3663 Affinity, rabbit) diluted in Immunol Staining Dilution Buffer (P0103, Beyotime) at 4 overnight. After incubation, cells were washed with PBS for 5 min each. Cells were then stained with IgG-Alexa 488 (1:500, AS035, ABclonal, Donkey) diluted in PBS with 10 μg/mL Hoechst 33342 at room temperature for 1 hr. Cells were then washed three times with PBS for 5 min each. Images were acquired, and process with an LSM 900 and further analyzed by ImageJ.

### Western Blot

Cells were lysed in a cold lysis buffer (50 mM Tris, pH 7.4, 150 mM NaCl, 1% Triton X100, 1% sodium deoxycholate, 0.1% SDS) supplemented with a protease/phosphatase inhibitor cocktail (78430, Thermo Fisher Scientific) on ice for 30 min. Protein concentration was quantified by Bradford method. The mitochondrial and nuclear proteins were extracted using Cell Nuclear Protein Extraction Kit (Proteintech, PK10014) and the Mitochondrial Extraction Kit (Proteintech, PK10016) following the manufacturer’s instructions. SDS-PAGE buffer was then added to same content of each sample, following denaturing at 95 for 5 min.

Equal amounts of total protein (30-50 μg per well) were separated by sodium dodecyl sulfate polyacrylamide gel electrophoresis and subsequently transferred onto a PVDF membrane. Membranes were then blocked with 5% milk or bovine serum albumen in TBST at RT for 2 hrs, and then washed three times with TBST for 10 min each. Membranes were then incubated overnight at 4 with the primary antibody (table S5), followed by three washes with TBST. The membrane was then incubated with the secondary antibody (table S5) at RT for 1 hr followed by a further three times of wash with TBST. The proteins were visualized using SuperSignal West Pico PLUS chemiluminescence ECL kit (34577, Thermo Fisher Scientific). Imaging was acquired using Bio-Rad ChemiDoc XRS+ (Biorad) and processed by ImageJ.

### Quantitative polymerase chain reaction analysis (Q-PCR)

Total RNA was isolated from HT-1080 cells utilizing RNAex Pro Reagent (AG21101 & AG21024, AG). The isolated RNA underwent reverse transcription by a two-step process to synthesize cDNA employing the EVO M-MLV RT Kit (AG11728, AG). Resultant cDNA was mixed with primer and 2× SYBR Green Pro Taq HS (AG11733, AG) and amplified using PCR amplifier (CFX Connect Real-Time System) The primer sequences were listed in table S5.

### Spatio-Temporal Mass Spectrometry in living cells

#### miniTurboID

HT1080 cells transiently transfected with miniTurboID plasmids were treated with 1.5 μM CuSO_4_ and 15 nM ES and harvested at a series of time points 0, 2, 4, 6, 8, and 10 hrs. 500 μM biotin (HY-B0511, MCE) was added into the medium 10 min prior to collection. After thoroughly washing with ice-cold PBS three times, cells were lysed in the lysis buffer (25 mM Tris, pH 7.2, 150 mM NaCl, 0.8% Triton X-100, and 1x protease inhibitor). Cell lysates were sonicated at 4, with three bursts of 10 s with an interval of 2 s between bursts. Cell debris was removed by centrifuged at 16,000 g for 20 min. The supernatant was then mixed with pre-washed Pierce™ Protein A/G Magnetic Agarose Beads (78609, Thermo Fisher Scientific) and gently agitated on a rocking platform at 4 for 2.5 hrs.

After incubation, the Streptavidin Agarose Resin was centrifuged at 500 g for 1 min, and the supernatant was removed. The resin was subjected to two washes with lysis buffer and four washes with PBS with each wash followed by centrifugation at 500 g for 1 min. 40 μL loading buffer (100 mM Tris-HCl, 4% SDS, 1 mM DTT, pH 7.6) was added to Streptavidin Agarose Resin, and the resin was then boiled at 100 for 20 min. After centrifugation at 14,000 g for 20 min to separate precipitation and supernatant, the supernatant was collected and stored at −80.

Protein quality was detected by Bradford protein quantitative method and 12% SDS-PAGE gel electrophoresis with coomassie brilliant blue staining. For quantitative proteomics studies, protein lysates were extracted using lysis buffer (8 M Urea, 100 mM TEAB, pH 8.5). The lysates were then reduced with 10 mM TCEP and alkylated with 50 mM iodoactamide, followed by acetone precipitation. The precipitated proteins were resuspended in 100mM TEAB and digested overnight at 37 with Trypsin (V5280, Promega). The resulting peptides are desalted using C18 desalting column (89852, Thermo Fisher Scientific. After three washes with a buffer containing 0.1% formic acid and 3% acetonitrile, the peptides were eluted with a buffer of 0.1% formic acid and 70% acetonitrile, and then lyophilized.

### Mass spectrometry workflow and data analysis

Mass spectrometry (MS)-based proteomics were analyzed by using the Parallel Accumulation Serial Fragmentation (PASEF®) technology with timsTOF Pro (Burker, America) at Novogene (China). 2D-LC-MS is performed on peptides, where in the first dimension of chromatographic analysis, SCX or high pH HPLC is primarily used to preliminarily separate the peptide mixture into multiple fractions. This step reduces the complexity of the peptide sample. Each of these fractions is then subjected to LC-MS to achieve better mass spectrometry identification results. Whole scanning range of MS was divided into several Windows, and all ions in each window were selected, broken and detected in a cycle.

The quantification type chosen is DIA (Data Independent Acquisition), Spectronaut-Pulsar is used to search the database and build the database. The parameters are: Mass tolerance for precursor ion was set to 10 parts per million (ppm), and for the product ion, it was set to 0.02 Daltons (Da). Carbamidomethylation of cystein was designated as fixed modification, Oxidation of methionine (M) was considered a variable modification. Acetylation was specified as a modification occurring at the N-terminus. Further screening using Spectronaut-Pulsar retained peptides with more than 99% confidence. Import the DIA data into the Spectronaut (Biognosys) software and extract ion pairs of chromatographic peaks. Calculate the peak area and achieve the qualitative and quantitative of peptides.

### Proteomics expression data preprocessing

Quantitative proteomics data preprocessing was conducted in R (v4.4.2) using a stratified quality control framework. For each organelle-time point combination (6 time points × 5 organelles), proteins were sequentially filtered based on the following criteria: (1) more than one missing value in the triplicate measurements; (2) detection in fewer than two non-zero biological replicates; (3) technical variability exceeding the 90th percentile of condition-specific coefficients of variation; and (4) mean abundance below the 10th percentile within their organelle-time group. After filtering, remaining missing values were imputed using condition-specific row means (excluding zeros when applicable). To achieve gene-level resolution, the most abundant protein isoform for each gene was retained. Filtered protein intensity values were then log2-transformed using a zero-preserving approach, where only non-zero values were transformed, while zero values were maintained as zeros. This transformation was performed independently for each organelle to preserve the biological significance of protein absence. Following transformation, datasets from individual organelles were merged into a comprehensive matrix. Zeros were preserved for proteins not detected in certain organelles. Finally, the mean intensity of each protein was calculated for each organelle–time point combination, resulting in a dataset of 2,867 high-confidence proteins with organelle-specific temporal profiles.

### Spatial-temporal proteomics expression pattern definition

To define spatiotemporal expression patterns, the preprocessed mean intensity of each protein was standardized using Z-score normalization. Clustering and heatmap visualization of the proteomics expression data were performed using the ComplexHeatmap package (v2.18.0), applying the Ward.D2 clustering method and Euclidean distance as the distance metric. Proteins were divided into ten clusters based on their expression patterns across organelles and time points by setting the row_km parameter to 10. For functional characterization, clusterProfiler (v4.10.0) was used to perform GO enrichment analysis on the genes within each cluster, with the background gene set restricted to GO Molecular Function. Pathways with p.adjust < 0.1 and FoldEnrichment > 1.5 were retained and sorted by FoldEnrichment. The resulting pathways were visualized to highlight the functional characteristics of each cluster, with the top 30 pathways shown for clusters enriched with more pathways.

### Differential protein expression analysis at each organelle

Differential expression analysis was conducted using the limma package (v3.60.6) on organelle-specific, zero-preserving log2-transformed data. For each organelle, protein abundance at experimental time points (2, 4, 6, 8, and 10 hours) was compared to the baseline (0 hours) using linear models with empirical Bayes moderation (trend=TRUE to account for intensity-dependent variance). Proteins with no detectable expression across all samples within a given organelle–time point comparison were excluded. Adjusted p-values were calculated using the Benjamini-Hochberg method to control the false discovery rate (FDR). Proteins were classified as differentially expressed if they met the thresholds of adjusted p-value < 0.01 and |log2 fold change| > 2. This analysis identified high-confidence differentially expressed proteins (DEPs) with distinct organelle-specific temporal expression patterns. Results were visualized using volcano plots, with color-coded markers highlighting significantly upregulated and downregulated proteins.

### Subcellular proteome trafficking analysis

In this analysis, organelle localization at each time point was determined using a maximum abundance-based method applied to the preprocessed quantitative proteomics data. Protein abundance data for each organelle and time point were extracted, and the coefficient of variation (CV) of replicate measurements was calculated. Proteins with a CV ≤ 0.3 were retained to ensure data reliability. Abundance values were then median-normalized across replicates to eliminate experimental biases. For each protein, normalized abundance values across organelles were ranked to identify the maximum abundance (Max_Abundance) and the second-highest abundance (Second_Max_Abundance). A confidence score was calculated as Confidence_Score = Max_Abundance / (Second_Max_Abundance + 0.01) to quantify the specificity of localization. Proteins were classified into three categories based on predefined thresholds: Unique (Confidence_Score > 10), Multiple (2 < Confidence_Score ≤ 10), and Uncertain (Confidence_Score ≤ 2). The organelle with the highest abundance was assigned as the final localization for each protein.

Proteins classified as Unique were further compared with known organelle annotations from publicly available databases, including Uniprot, The Human Protein Atlas, and Gene Ontology Cellular Component (GO CC), to evaluate the consistency of predicted localizations with existing knowledge. The distribution of protein assignment types (Unique, Multiple, and Uncertain) was summarized and visualized across all time points to assess temporal dynamics in organelle localization. To illustrate subcellular trafficking patterns, the movement of proteins among five organelles across six time points was visualized using a Sankey diagram generated with the ggalluvial package (v0.12.5) and ggplot2 package (v3.5.1).

### Statistical Analysis

Statistical analysis was performed using GraphPad Prism (8.3.1). All experimental data shown in plots are represented as mean ± SD/SEM, and exact number of independent replicates for each experiment is stated in figure legends. Significant differences were determined by one-way ANOVA or two-tailed t-tests. The significance level between two groups was set to 0.05 and the exact *p*-values were shown in the figures. For conjugation frequency experiments, analyses were performed on log-transformed data.

## Supporting information

tableS1-filtered_protein_abundance

tableS2-DEP_all

tableS3-localization_assignment

Table S4-The boutique RNAi and overexpression screen

Table S5-List of consumables and primers

## Acknowledgments

This research was funded by the Zhejiang Provincial Natural Science Foundation of China (No. LD22H310004), National Natural Science Foundation of China (No. 82473894), CAMS Innovation Fund for Medical Sciences (CIFMS, 2019-I2M-5-074) to C, F; National Natural Science Foundation of China (No.82073412) to Q, Z; J. K received support from the 5Point Foundation (Christine Martin Fellowship).

## Author contributions

Conceptualization: CF, QZ and JK.

Methodology: CF, QZ and XY.

Investigation: XY, CW, GS, SZ, KZ and XD.

Formal analysis: CS and XL.

Visualization: XY and CW.

Funding acquisition: CF, QZ and JK.

Supervision: CF.

Writing – original draft: QZ.

Writing – review & editing: JK, CF, CS.

---

**Extended Data Fig. 1.**
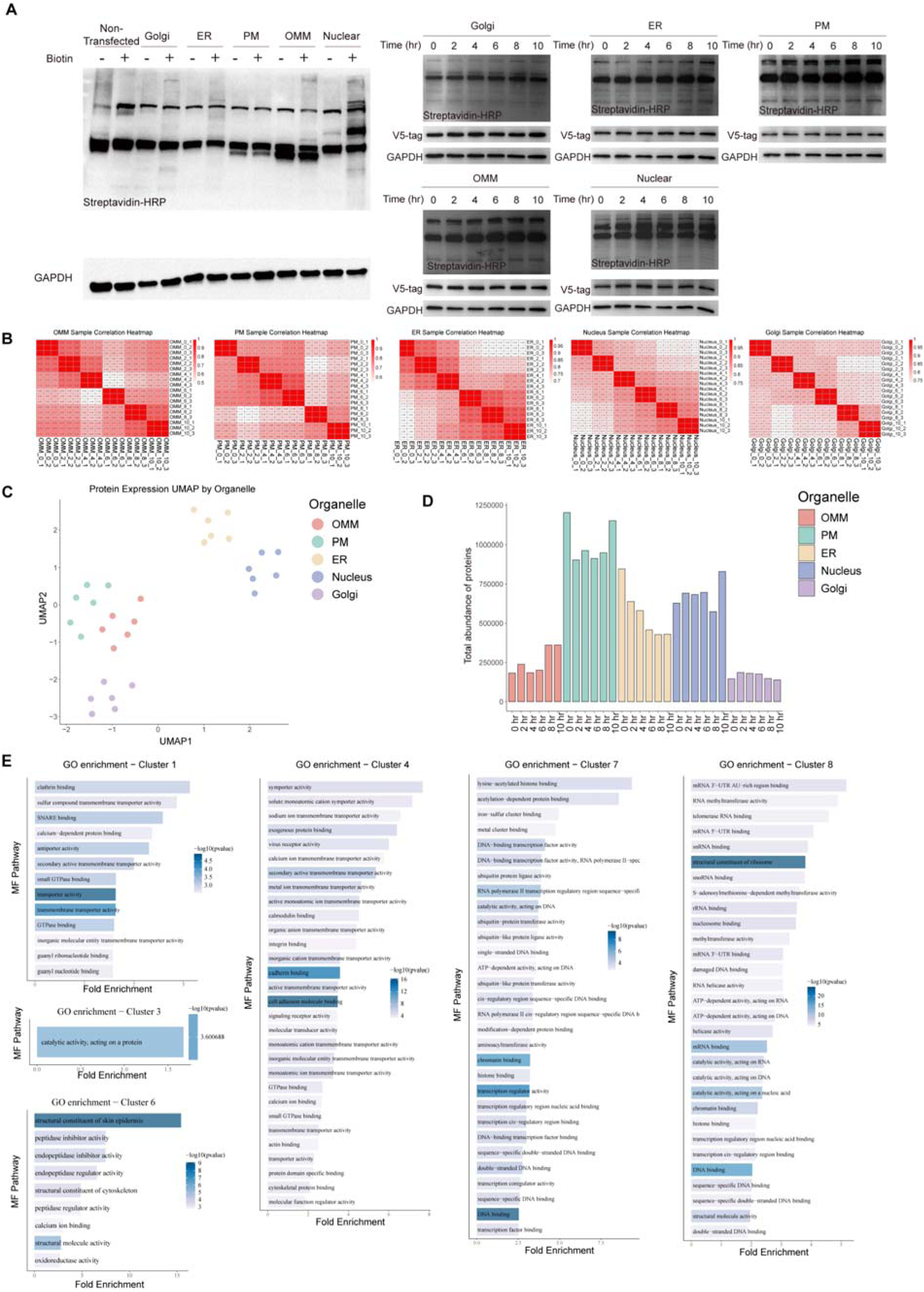
Spatio-Temporal Mass Spectrometry analysis delineates subcellular proteome dynamics in early cuproptosis. (**A**) Western blot analysis of biotinylated proteins using HRP-conjugated Streptavidin and biotin ligase expression by anti-V5 in different organelles of cells transfected with miniTurboID. Cells were labelled with 500 μM exogenous biotin after exposure to 1.5 μM CuSO_4_ and 15 nM ES at the indicated time points. (**B**) Analysis of the correlation among replicate samples of Spatio-Temporal Mass Spectrometry. (**C**) Uniform manifold approximation and projection (UMAP) visualization of protein expression data in five organelles across six time points. (**D**) Total protein abundance in each organelle at different time points. (**E**) Enrichment analysis of genes in each cluster identified in Fig. 1E using GO molecular function gene set with clusterProfiler (No pathways were retained for cluster 2 and 9 due to failing to meet the threshold of p.adjust < 0.1 and FoldEnrichment > 1.5).

**Extended Data Fig. 2.**
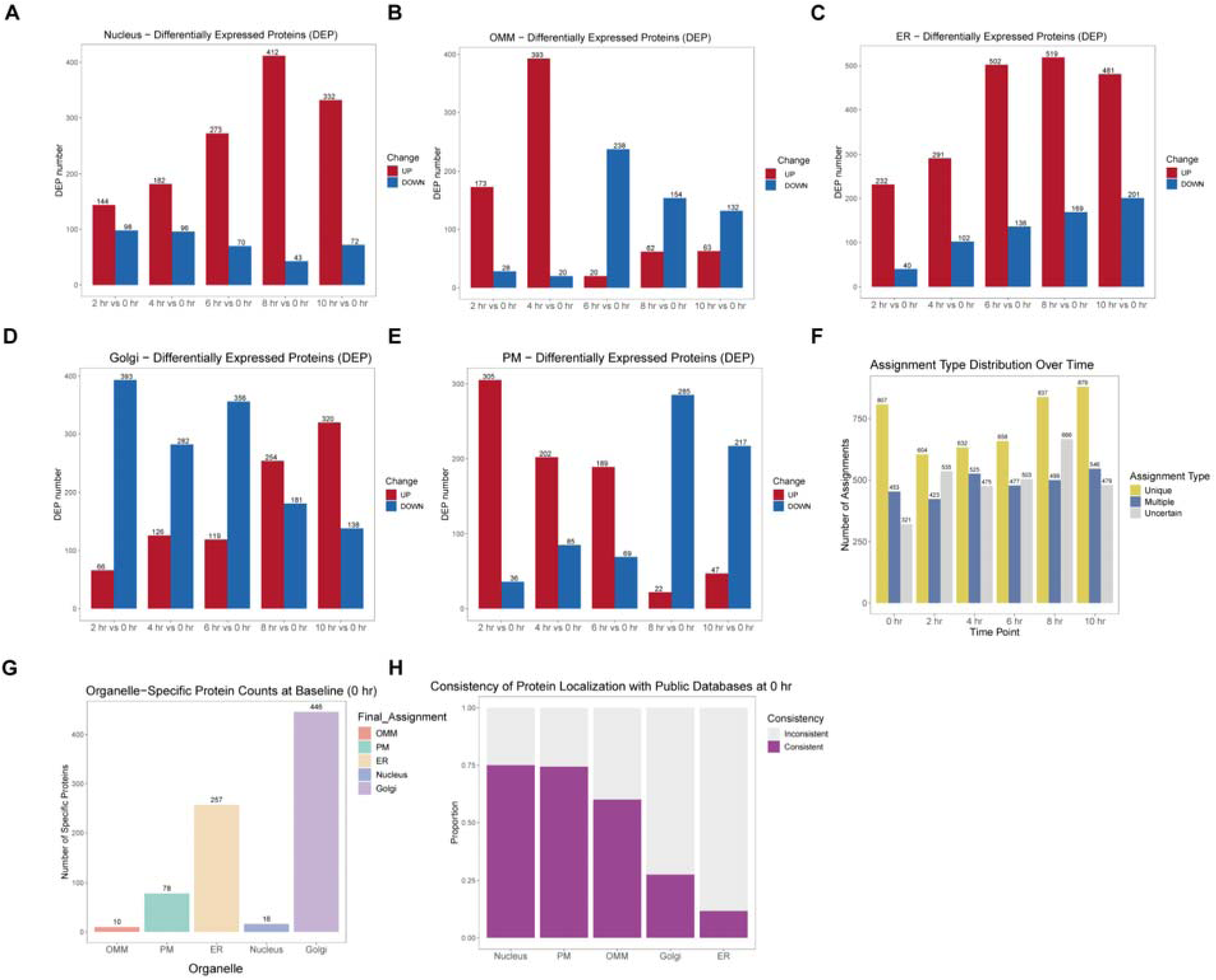
Spatio-Temporal Mass Spectrometry analysis identifies differentially expressed protein across various organelles over time. (**A**-**E**) The number of differentially expressed proteins identified in the nucleus (A), outer mitochondrial membrane (OMM) (B), ER (C), plasma membrane (PM) (D) and Golgi (E) at 2, 4, 6, 8 and 10 hrs compared to 0 hr. **(F)** The distribution of protein assignment types (“Unique”, “Multiple” and “Uncertain”) across different time points. **(G)** The number of proteins categorized as “Unique” across organelles at baseline (0 hr). **(H)** The proportion of proteins (categorized as “Unique”) whose localization is consistent or inconsistent with public database annotations at baseline.

**Extended Data Fig. 3.**
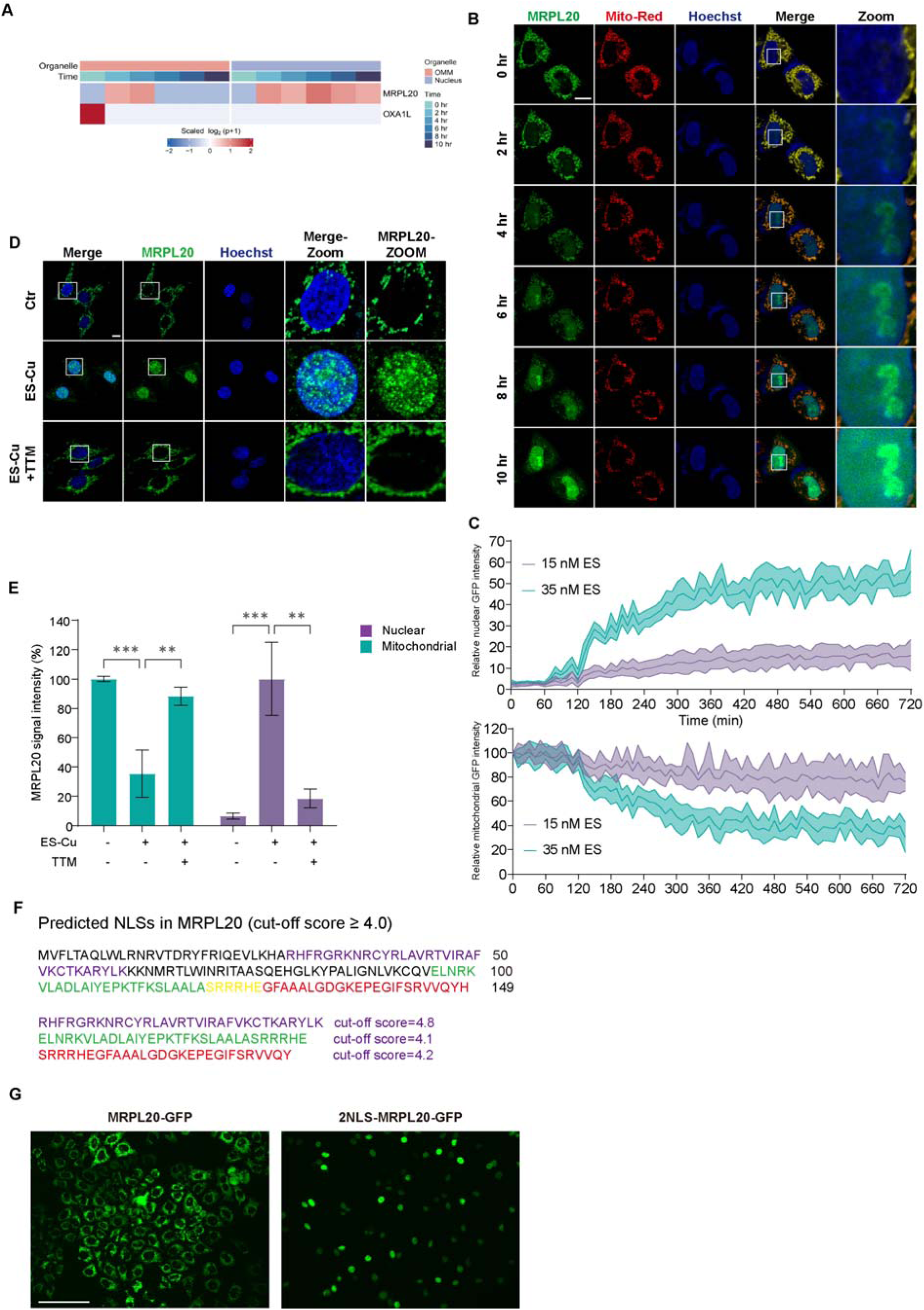
Mitochondria-nucleus shuttling of MRPL20 is a cuproptosis specific event. (**A**) Spatio-Temporal Mass Spectrometry analysis of MRPL20 and OXA1L expression in outer mitochondrial membrane (OMM) and nucleus at six time points upon treatment with ES-Cu. (**B**) Time-lapse fluorescence images showing mitochondrial and nuclear MRPL20-GFP expression in HT-1080 cells. Cells overexpressing GFP-tagged MRPL20 were treated with 1.5 μM CuSO_4_ and 35 nM ES and subject to image analysis at the indicated time points. Scale bars, 10 µm. (**C**) Histograms showing quantification of MRPL20-GFP intensity normalized to mitochondrial GFP intensity at time 0. Data are presented as mean ± SD, n = 6. (**D**) Fluorescence images showing endogenous MRPL20 expression in mitochondria and nuclei of HT-1080 cells. Cells were treated with 1.5 μM CuSO_4_ and 15 nM ES in the presence or absence of 20 μM TTM for 12 hrs before being fixed and stained using anti-MRPL20 FITC antibody. Scale bars, 10 µm. (**E**) Histograms showing quantification of endogenous MRPL20 expression. Data are presented as mean ± SEM, n = 6. Statistical analysis was performed using a one-way ANOVA. (**F**) Predicated intrinsic nuclear-localization signal (NLS) in MRPL20 using the cNLS Mapper tool. (G) Fluorescence images of MRPL20-GFP and 2NLS-MRPL20-GFP (MRPL20 fused with an exogenous NLS) expression in HT-1080 cells. Scale bars, 125 µm. * *p*-value < 0.05, ** *p*-value < 0.01, *** *p*-value < 0.001, **** *p*-value < 0.0001.

**Extended Data Fig. 4.**
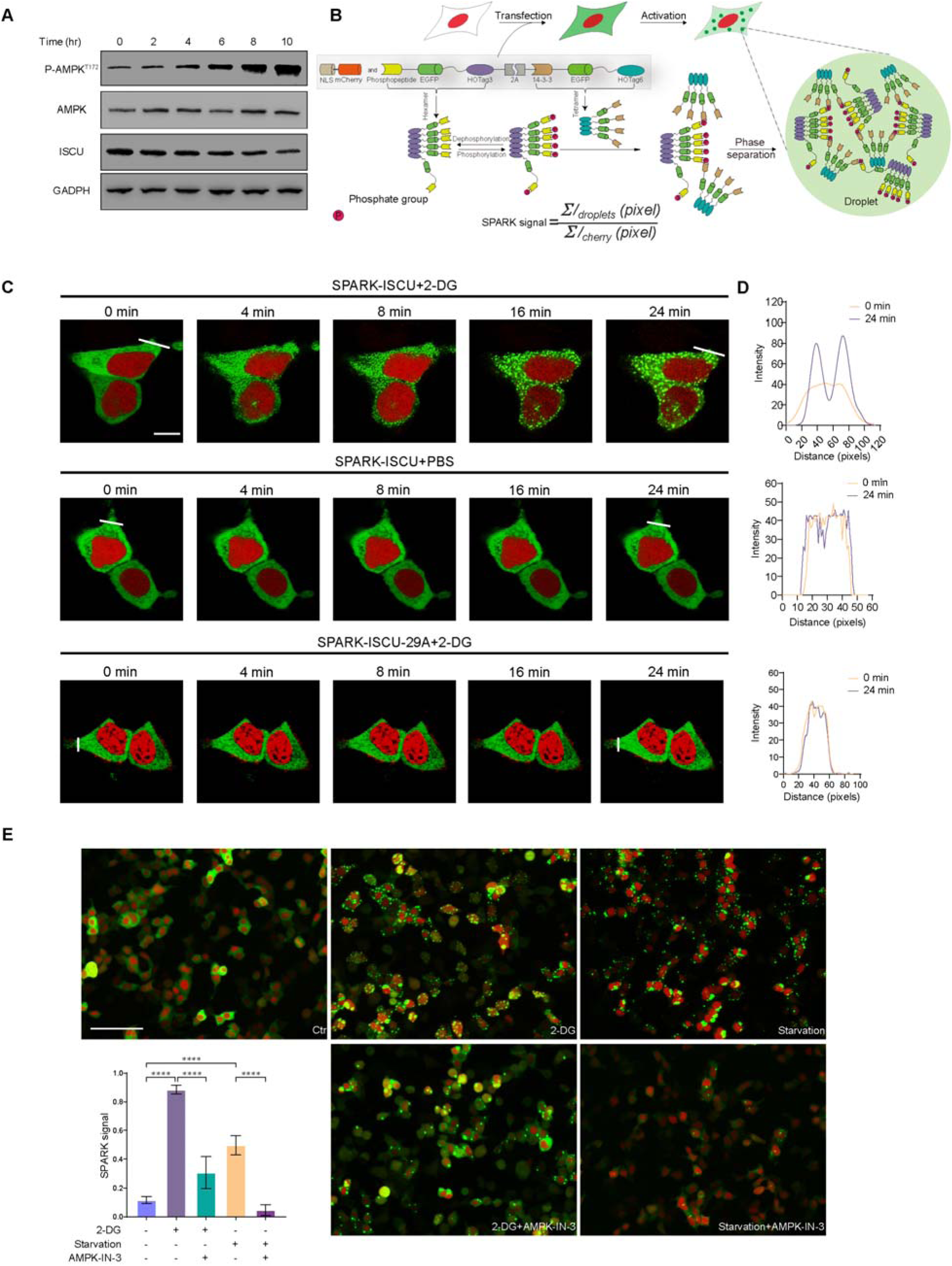
Visualizing AMPK activity in living cells with AMPK-SPARK during cuproptosis. (**A**) Western blot analysis of AMPK-ISCU signaling pathway. HT-1080 cells were treated with 1.5 μM CuSO_4_ and 15 nM ES for different time periods as indicated. GAPDH is used as a loading control. (**B**) Schematic showing the principle of AMPK-SPARK. HOTag3 can oligomerize into hexamers and HOTag6 into tetramer. AMPK Substrate peptide of ISCU can be recognized by phosphoserine binding domain of 14-3-3 after phosphorylation. Cells were transiently co-transfected with AMPK-SPARK plasmid and NLS-mCherry plasmid. To quantitatively measure the droplets assembly/disassembly over time, we define SPARK signal as the sum of GFP fluorescent droplets’ pixel intensity divided by sum of cells’ cherry pixel intensity. (**C**) Time-lapse fluorescence images of cells expressing AMPK-SPARK or the mutated reporter that cannot be phosphorylated by AMPK upon treatment with 50 μM 2-DG. Scale bars, 10 µm. (**D**) Charts corresponded to the images. (**E**) Fluorescence images of HEK293T cells expressing AMPK-SPARK upon different treatments. Cells were treated with 50 μM 2-DG or cultured in PBS for 30 min with or without pre-treatment of 25 μM AMPK-IN-3 for 2 hrs. Scale bars, 125 µm. Data are presented as mean ± SD, n = 3. Statistical analysis was performed using a one-way ANOVA. * *p*-value < 0.05, ** *p*-value < 0.01, *** *p*-value < 0.001, **** *p*-value < 0.0001.

**Extended Data Fig. 5.**
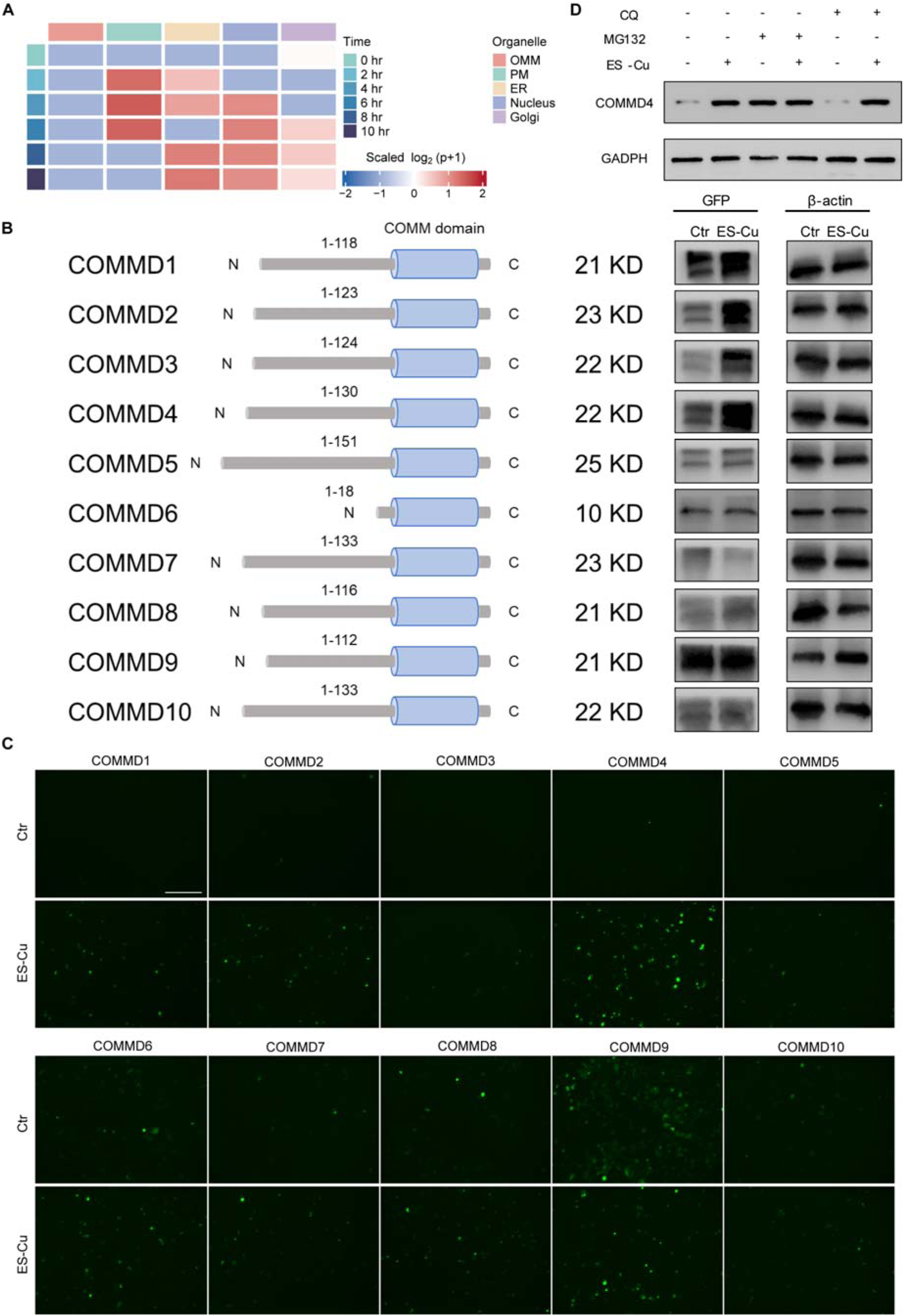
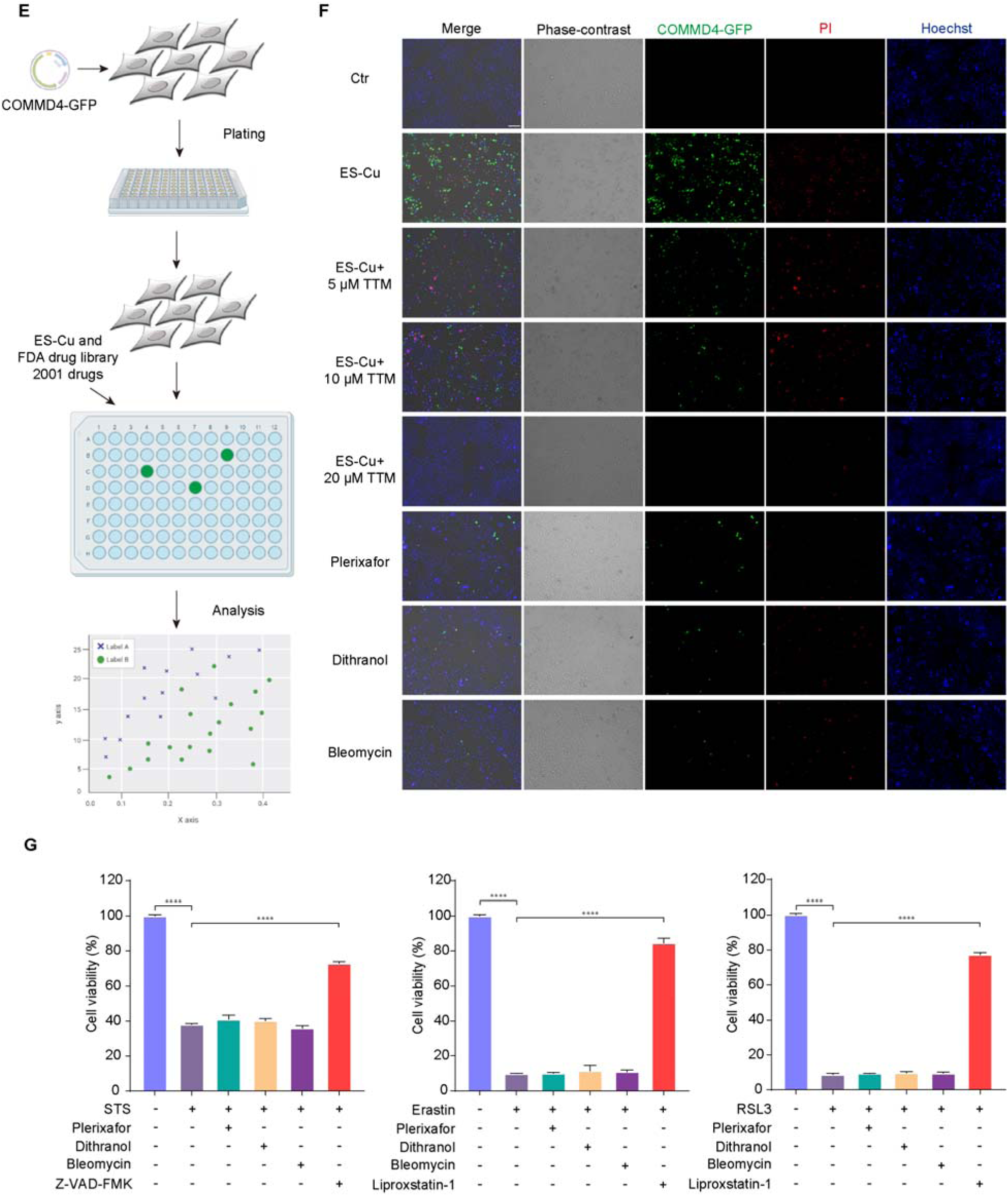
COMMD4 is a is a biomarker of cuproptosis. (**A**) Spatio-Temporal Mass Spectrometry analysis of COMMD4 expression in different cellular compartments at six time points upon treatment with ES-Cu. (B) Schematic of structure of the COMMD family members (left) and western blot images showing expression of GFP-tagged COMMD family members upon treatment with 1.5 μM CuSO_4_ and 15 nM ES for 12 hrs (right). β-actin is used as a loading control. (**C**) Fluorescence images of HT-1080 cells transfected with GFP-tagged COMMD family members after treatment with 1.5 μM CuSO_4_ and 15 nM ES for 12 hrs. Scale bars, 275 µm. (**D**) Western blot images of COMMD4 expression in HT-1080 cells pre-treated with 10 μM MG132 or 50 μM CQ followed by 1.5 μM CuSO_4_ and 15 nM ES for 24 hrs. (**E**) Schematic diagram of the working process of high-content screening for the FDA approved drugs library. (**F**) Fluorescence images of Hela cells transiently transfected with GFP-tagged COMMD4 after treatment with Plerixafor, Dithranol, and Bleomycin at 5 μM in combination with 1.5 μM CuSO_4_ and 35 nM ES for 24 hrs. Scale bars, 275 µm. (**G**) Viability of HT-1080 cells pretreated 2 hrs with either 5 μM Plerixafor, 5 μM Bleomycin, 5 μM Dithranol, 10 μM Z-VAD-FMK or 1 μM liprozstatin-1 followed by treatment with either 1 μM STS, 5 μM Erastin or 250 nM RSL3 for 24 hrs. Data are presented as mean ± SD, n = 3. Data are presented as mean ± SD, n = 3. Statistical analysis was performed using a one-way ANOVA (F), (G) and (H). * *p*-value < 0.05, ** *p*-value < 0.01, *** *p*-value < 0.001, **** *p*-value < 0.0001.

